# Spatiotemporal Proteome Remodeling Directs Human Hematopoietic Stem Cells via a Metabolic-Cytoskeletal Axis

**DOI:** 10.64898/2026.06.08.728414

**Authors:** Zahra Masoumi, Navin Shirodkar, Lauren Wheeler, Megan Guthrie, Daniel M. Davies, Diana Hernandez, William Grey

**Affiliations:** ProteoStem Lab, Centre for Blood Research, York Biomedical Research Institute, Department of Biology, University of York, UK; Anthony Nolan Research Institute, Royal Free Campus, London, UK; UCL Cancer Institute, Royal Free Campus, London, UK

## Abstract

Hematopoietic stem cells (HSCs) sustain the lifelong production of all blood and immune cells and are central to transplantation and gene therapies. However, the clinical use of these approaches remains limited due to the functional attrition of HSCs during *ex vivo* adaptation and expansion in culture. As these transitions are executed at the protein and structural level, the primary drivers of cellular adaptation have remained elusive in transcriptional studies. Here, we overcome the technical resolution limits of rare primary cells by integrating a scalable, ultra-low input proteomics pipeline with 3D super-resolution imaging to map the proteomic and spatial organization of primary human HSCs during *ex vivo* expansion. Our analysis reveals a previously unrecognized, metabolic-cytoskeletal axis that coordinates organelle reorganization and structural polarity. Perturbation of this axis preserves structural polarity while uncoupling it from biomass expansion, demonstrating that growth and functional deterioration are separable processes in *ex vivo* HSC culture. These findings establish a framework for understanding *ex vivo* HSC adaptation and expansion and inform the rational development of next-generation cellular therapies.

## Introduction

Hematopoietic stem cells (HSCs) sustain the lifelong production of all blood and immune cells through a balance of self-renewal and multilineage differentiation. This capacity is the foundation of hematopoietic homeostasis and the clinical success of hematopoietic cell transplantation (HCT), a curative therapy for a wide range of hematological malignancies, immunodeficiencies and emerging gene-based treatments^1,2^. However, the broader application of HCT remains limited, particularly due to progressive loss of HSCs and their function^3,4^ during *ex vivo* adaption^5^ and expansion^6,7^. A mechanistic understanding of the molecular pathways governing *ex vivo* adaptation is, therefore, paramount to moving beyond empirical optimizations^8–10^ and rationally improving the efficacy of future culture systems.

Current insights into HSC identity and function during *ex vivo* expansion rely heavily on single-cell transcriptomics^5,11,12^. However, the critical drivers of cellular adaptation and expansion including metabolic rewiring, cytoskeletal remodeling, and organelle dynamics are executed at the protein level, and RNA abundance is often a poor proxy for functional state. Despite significant advances in the proteomics field^13–17^, the landscape of human HSCs remains poorly defined at protein level, due to a combination of technical difficulties, low coverage, high cost, or time-intensive workflows that are currently available for analyzing low protein inputs from small primary cells in rare populations.

Equally, we lack a clear understanding of how proteomic remodeling translates into physical cellular architecture. While RAS and RHO GTPases are known to orchestrate cytoskeletal networks, organelle inheritance, and intracellular polarity that governs stem cell fate decisions^18–24^, resolving these spatial features in HSCs has remained a challenge in the field. The in-suspension nature of HSCs complicates high-resolution imaging and often requires artificial tethering strategies^19,20^ that inadvertently perturb the intrinsic polarity, organelle dynamics or signaling programs under investigation. Consequently, the structural process of HSC adaptation remains elusive, leaving a gap between the molecular picture and the physical execution in the cell.

Here, we integrate a scalable, ultra-low input proteomics pipeline with a 3D super-resolution imaging workflow, specifically tailored for suspension cells, to map the coordinated molecular and functional landscape of human HSCs, *de novo* and during *ex vivo* culture. Our analysis reveals a previously unrecognized metabolic-cytoskeletal axis that governs organelle reorganization and HSC adaptation. Functional perturbation of this axis, using independent inhibitors of fatty acid synthase and tubulin polymerization, demonstrated the significance of the metabolic-cytoskeletal coupling in regulating early HSC adaptation and subsequent expansion trajectory in culture. This work generates the first spatiotemporal proteomic and functional roadmap of *ex vivo* HSCs in culture and establishes a biology-first foundation for future expansion strategies and cell-based therapies.

## Results

### A robust ultra-low input proteomics platform reveals proteome-wide remodeling of HSCs in culture

To overcome the depth and throughput limitations of current proteomic methodologies, we systematically evaluated and optimized the lower limits of a DDM-based one-pot lysis protocol for rare, sorted HSC populations with low protein content. Using a Bruker TimsTOF HT, we demonstrated comprehensive proteome coverage capturing full subcellular localization (Supplementary Figure 1). We applied this methodology to investigate proteomic dynamics during *ex vivo* expansion of human HSCs. Ten pools of cord blood CD34^⁺^ cells from 30 donors were cultured in a clinically compliant culture system^25–28^, with or without UM729, for up to 7 days. On days 0 (*de novo*), 1 (adaptation), 3 (cell cycle progression), and 7 (proliferation/exhaustion), 200 immunophenotypic HSCs (see Methods) were sorted for proteomic analysis (Figure 1a; Supplementary Figure 2a). Total cell numbers decreased between day 0 and 1 and subsequently increased throughout culture (Figure 1b; Supplementary Figure 2b). Immunophenotypic long term HSCs declined overnight in cytokine only conditions, with modest proliferation between days 3 and 7. On the other hand, UM729 treatment stabilized HSC numbers initially (in the majority of pools) and promoted expansion across all timepoints (Figure 1b; Supplementary Figure 2b).

**Figure 1.**
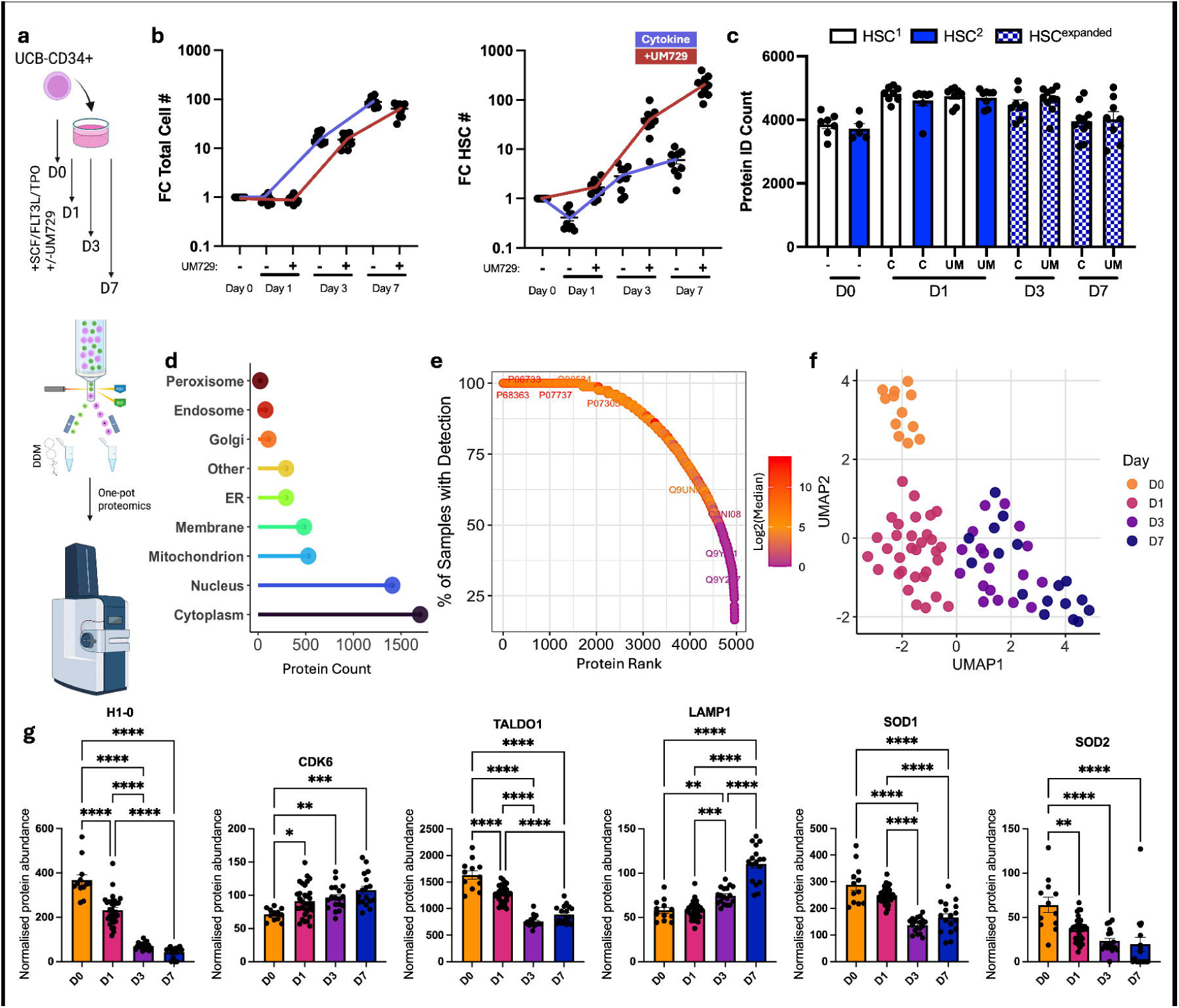
Proteome-wide remodeling of HSCs during adaptation to culture. **a**, Schematic of cell culture and ultra-low input proteomics pipeline, **b**, Fold change in total and phenotypic HSC cell counts (#) vs day 0 input on days 1, 3 and 7 of culture (+/- UM729), **c**, Protein ID counts from proteomics on sorted phenotypic HSCs per day in culture (C = cytokines only, UM = +UM729, HSC^1^: CD49f^+^ EPCR^+^, HSC^2^: CD49f^+^ EPCR^-^, expanded HSC^e^: CD90^+^ EPCR^+^), **d**, The number of proteins detected based on their cellular localizations, **e**, Data completeness ranked based on mean expression of protein across the dataset, **f**, UMAP of the HSC proteome colored by days in culture, **g**, Comparison of normalized abundance of H1-0, CDK6, TALDO1, LAMP1, SOD1 and SOD2 in HSCs grouped by days in culture (one-way ANOVA was used to calculate statistical significance, * = p<0.05, ** = p<0.005, *** = p<0.0005, **** = p<0.0001).

Global proteome analysis of HSCs resulted in identification of a total of 5765 protein groups across all conditions and all major subcellular compartments (minimum 2 unique peptides per protein, FDR < 0.01, Fig. 1c, d). From the 5621 protein groups differentially abundant in at least one comparison, a core proteome of 3534 proteins was quantified in ≥80% of samples, reflecting high data completeness and low protein dropout (Figure 1e, Supplementary Figure 2c). The high coverage and consistency in our data completeness enabled robust high-resolution comparative and trajectory-based analyses, without the need for alternative foundational data or shared latent space from transcriptomics^17^. Unbiased PCA and UMAP representation of the data revealed distinct clustering as HSCs transitioned through culture (Figure 1f; Supplementary Figure 3a). While the overall proteomic similarity remained high (63%), reflecting the preservation of cell identity, temporal progression induced significant changes in global proteomic profiles of HSCs (Figure 1f). Crucially, these shifts reflected remodeling of the proteomic profile and were not skewed by the over-or under-representation of specific subcellular compartments (Supplementary Figure 3b), and included significant shifts in markers of quiescence or activity (e.g. H1-0 and CDK6)^29,30^ as well as established functional regulators (e.g. SOD1, SOD2, LAMP1 and TALDO1) of HSCs^17,19^ (Figure 1g). Interestingly, the proteomics data captured a transition from “LT-HSC up” to “LT-HSC down” signatures, originally defined by transcriptomics^31^ (Supplementary Figure 3c). Together, these results demonstrate a reproducible, high-depth roadmap of the proteomic shifts underpinning the adaptation and proliferation of human HSCs in culture.

### Proteomic trajectories identify a metabolic-structural axis during *ex vivo* HSC adaptation and expansion

To define the progressive proteomic states of HSCs during *ex vivo* culture, we applied unsupervised pseudotime reconstruction and trajectory analysis to EPCR^+^ HSCs and compared the outcome with supervised analyses based on experimental timepoints. The unsupervised analysis recapitulated the temporal progression observed across culture timepoints (Figure 1f), revealing five distinct temporal transition modules (Figure 2a, b). Signatures of oxidative phosphorylation, MYC targets and RNA processing and splicing were present in two modules: proteins in Module 1 were most strongly present in day 0 and day 7 HSCs, and proteins in Module 2 were maintained through adaptation but gradually decreased and were lowest by day 7 (Figure 2c, Supplementary Figure 4a). Additionally, Module 1 contained a fatty acid metabolism signature that was upregulated in both day 0 and day 7 HSCs. Signatures in Module 3, associated with mTORC1 signaling, E2F targets, G2/M checkpoint, protein translation and unfolded protein response, were progressively obtained as cells adapted to culture and transitioned toward proliferation by day 3 and 7 (Figure 2c, Supplementary Figure 4a). However, certain MYC targets and mTORC1 signaling-associated proteins were most highly expressed in day 7 HSCs as shown in Module 5 (Figure 2c). To validate these findings, we performed supervised differential expression pattern analysis using the known time in culture, which revealed HSC proteomic profiles across the culture period (Supplementary Figure 4b). Importantly, proteins from the Tradeseq-derived temporal modules exhibited concordant expression dynamics across the experimental time course (Figure 2d), confirming that unsupervised analysis accurately captured the underlying temporal proteomic transitions, demonstrating the resolution and capacity of our dataset for unbiased trajectory inference.

**Figure 2.**
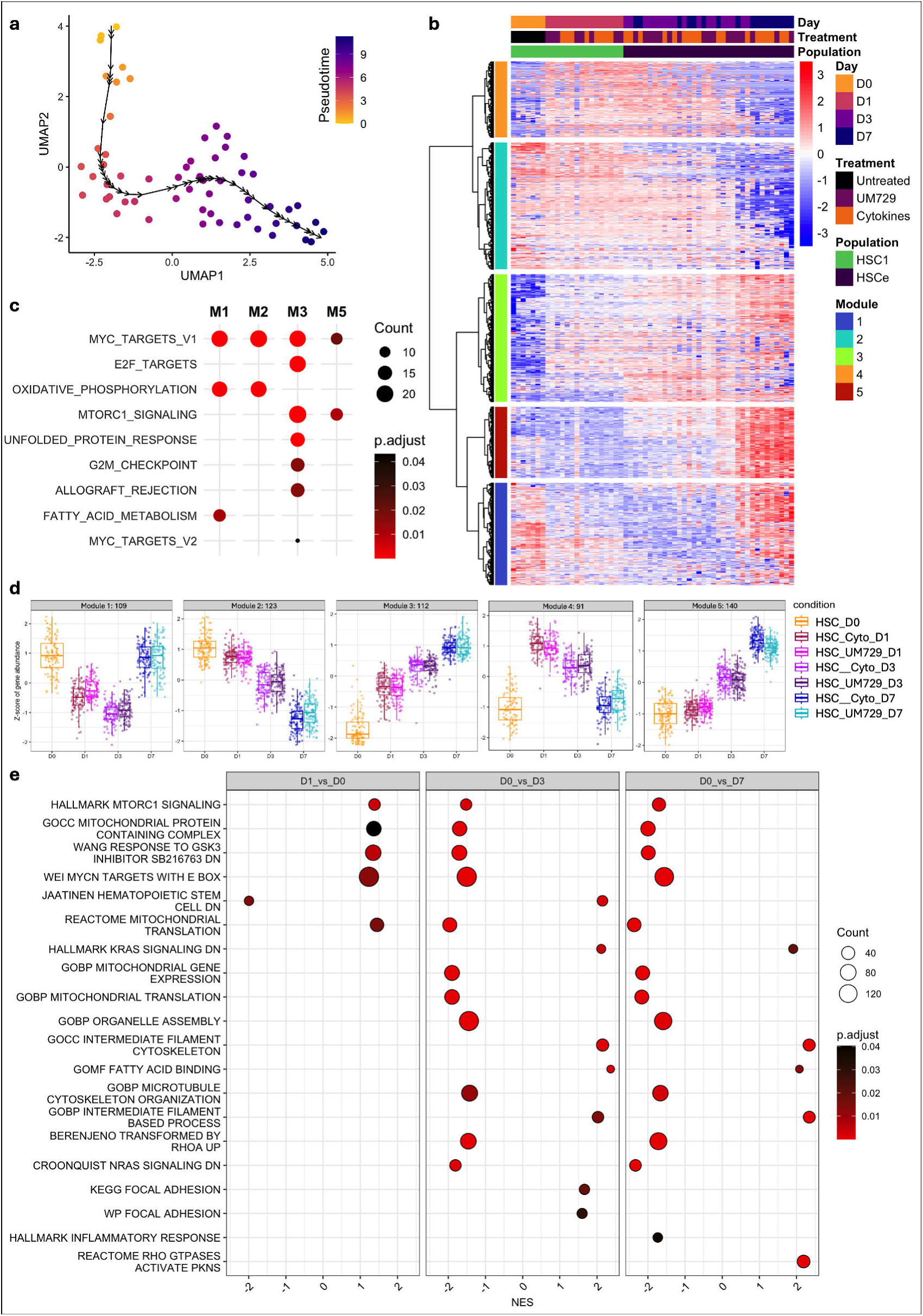
Dynamic signaling, metabolic, and structural transitions in HSCs during adaptation and expansion in culture. **a**, Pseudotime analysis of the total HSC proteome colored by pseudotime, **b**, Heatmap showing z-scores of significantly differentially abundant proteins across the pseudotime using Tradeseq, **c,** Dot plot showing the enriched Hallmark pathways associated with each module from Tradeseq analysis changing across pseudotime, **d,** DEGpatterns showing supervised temporal expression changes of proteins from each Tradeseq module, **e**, GSEA analysis of the indicated comparisons from total proteomics during adaptation to and expansion in culture, colored by p.adj value, showing selected significant pathways.

To validate the temporal and trajectory analyses, we further examined the transition states through gene set enrichment analysis (GSEA). The initial cytokine stimulation (day 1) was marked by a rapid induction of mTORC1 signaling, that was sustained through day 7, alongside MYC targets, and a drop in hematopoietic stem cell signature (Figure 2e). Day 3 was characterized by a coordinated enrichment of G2/M checkpoints (Figure 2e, Supplementary Figure 4c). By day 7, G2/M checkpoints were present along with an enriched E2F-target (Supplementary Figure 4c) suggesting progress in proliferative state. Concurrent with these changes were unique RAS signaling, cytoskeletal and metabolic signatures at days 0, 1, 3 or 7 (Figure 2e, Supplementary Figure 4c). Together, these data suggest that HSC expansion is not simply a proliferative switch, but rather involves a coordinated transitional state characterized by changes in metabolic flux and cytoskeletal dynamics.

### Coordinated shifts in FASN and RAS signaling reflects a coupled metabolic-cytoskeletal landscape in HSCs

To identify the regulators upstream of the signaling and structural changes that govern HSC adaptation and expansion, we examined proteins exhibiting rapid and sustained induction following cytokine exposure. Fatty acid synthase (FASN) showed a sharp increase within the first 24 hours of culture, with or without UM729, a trend that persisted through day 7 (Figure 3a, b, Supplementary Figure 4d). Interestingly, FASN induction was synchronized with shifts in abundances of key structural regulators, including the Ras GTPase-activator IQGAP2, the microtubule stabilizer TPT1 and the mitochondrial enzyme SHMT2 (Figure 3a, b).

**Figure 3.**
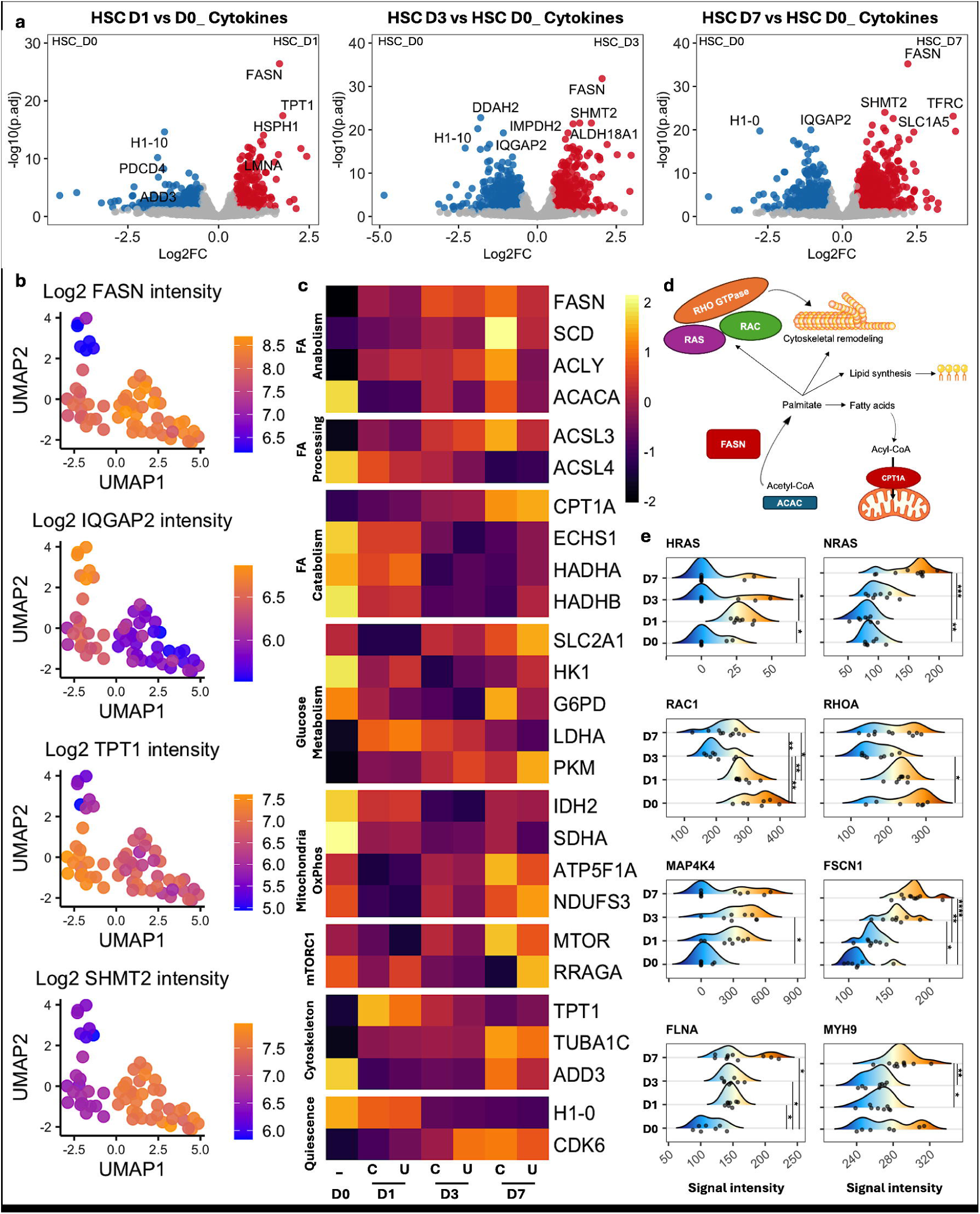
Coordinated FASN activation and RAS downregulation orchestrate metabolic and cytoskeletal remodeling in HSCs. **a**, Volcano plots of total proteomics (no imputation) in HSCs during adaptation to culture comparing day 0 HSCs to days 1, 2 and 7 (red = log2FC >0.5 and p.adj <0.05, blue = log2fc <-0.5 and p.adj <0.05) **b**, Log2-transformed abundance of selected proteins overlaid on the UMAP showing temporal changes in culture, **c**, Heatmap of scaled abundance of proteins activated or modified via FASN, regulating metabolic and cytoskeletal remodeling in HSCs as they adapt to culture or expand in basal media (C) or +UM729 (UM), **d**, Schematic representation of FASN-driven regulation of lipid biosynthesis and cytoskeletal remodeling, **e**, Ridge plots showing changes in RAS/RHO GTPase-driven signaling and cytoskeletal proteins in HSCs across days in culture (Dunn’s nonparametric pairwise comparison, following a statistically significant Kruskal Wallis (p value < 0.05), was used to calculate statistical significance, * = p<0.05, ** = p<0.005, *** = p<0.0005, **** = p<0.0001).

Further proteomic analysis revealed a comprehensive metabolic switch, specifically an early transition from catabolic FAO state (e.g., ECHS1, HADHA) toward active FA biosynthesis (e.g., FASN, ACLY, SCD) (Figure 3b, c). Critically, this metabolic switch was coordinated with microtubular and cytoskeletal remodeling (e.g., TPT1, ADD3) within the first 24 hours as cells adapt to culture, prior to the onset of proliferation. (Figure 3b, c). Given that FASN-generated palmitate is a primary substrate for both membrane expansion and palmitoylation of proteins such as tubulin and RHO-family effectors^32–35^, we further interrogated RAS and RHO GTPase signaling as well as cytoskeletal dynamics (Figure 3d). Early adaptation, within the first 24 hours, was characterized by a coordinated reduction in RAC1 and RHOA, alongside transient HRAS activation and MAP4K4 induction that was sustained through day 7 (Figure 3e). Coupled with increased FASN expression, these early shifts likely facilitate an early restructuring of microtubule and cytoskeletal architecture. This early adaptation phase was followed by progressive increases in FSCN1 and FLNA and suppression of RAC1/RHOA, reflecting further cytoskeletal remodeling (Figure 3d, e). By day 7, NRAS upregulation and bimodal RHOA distribution supported the emergence of altered biomechanical profiles (Figure 3d). Together, these proteomic signatures suggest a new metabolic-cytoskeletal axis, where early FASN-mediated lipid biosynthesis is coupled to RAS-regulated cytoskeletal dynamics and governs *ex vivo* adaptation of HSCs prior to expansion.

### Super resolution imaging reveals temporally distinct cytoskeletal and mitochondrial remodeling during HSPC adaptation

To determine whether the temporally dynamic proteomic changes translated into spatial remodeling of cellular architecture, we established a customized super-resolution imaging workflow optimized for suspension cells, enabling quantitative visualization of cytoskeletal and organelle organization in their native 3D state (Figure 4a, Supplementary Figure 5; see Methods). Following imaging and volumetric reconstruction of HSPCs across culture timepoints (Figure 4b), structural segmentation and quantitative analysis was performed (Supplementary Figure 5), enabling robust extraction of morphological features including volumetric and surface parameters for microtubule filaments and mitochondria (Figure 4c).

**Figure 4.**
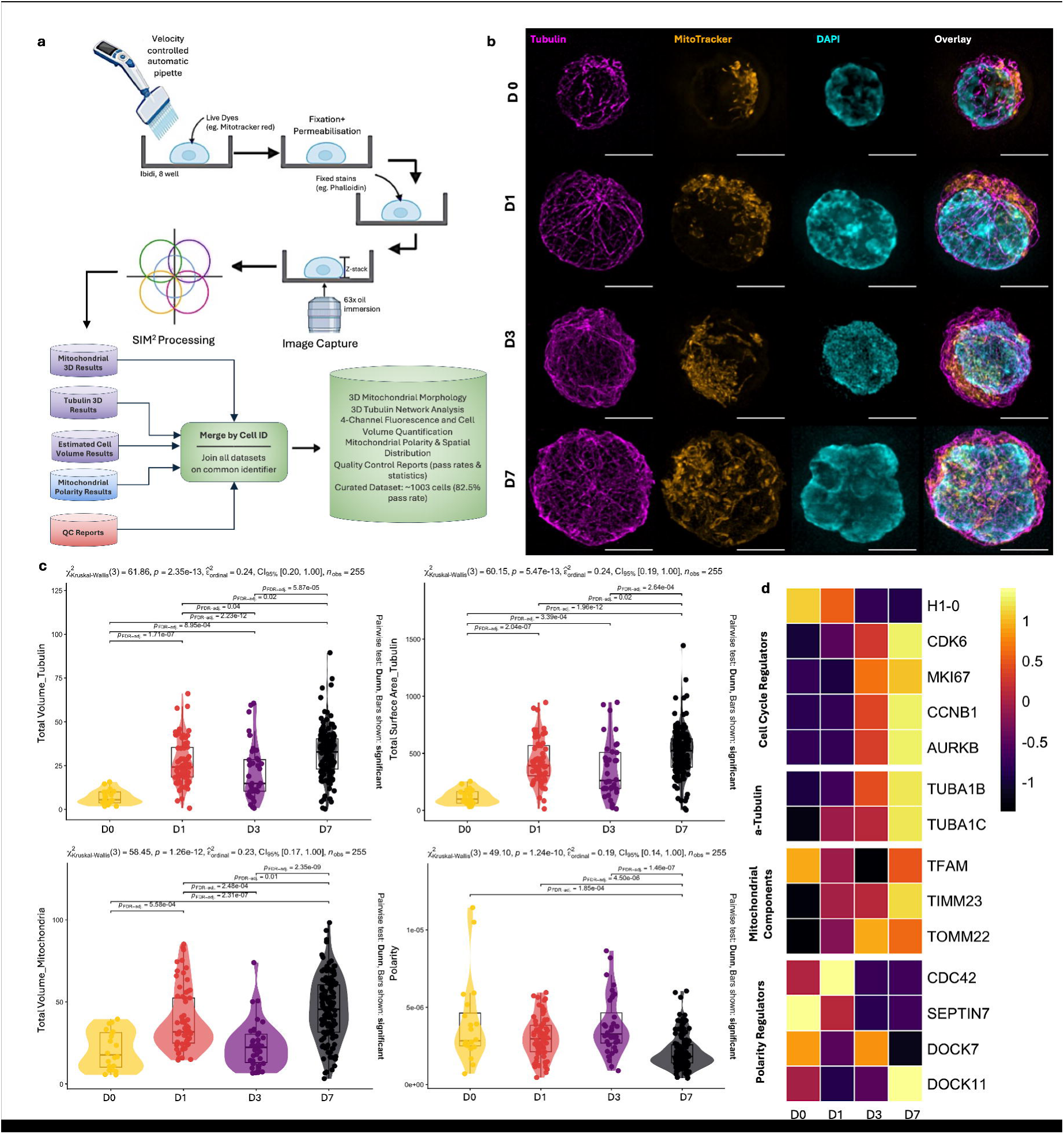
Spatial imaging of temporal cytoskeletal and mitochondrial dynamics during HSPC adaptation and expansion in culture. **a,** Schematic of sample preparation and microscopy technique used for super-resolution imaging of HSPCs, **b**, Representative images of single cells across days 0, 1, 3 and 7 (from top to bottom) in culture showing tubulin (magenta), mitochondria (MitoTracker Red, orange), nucleus (DAPI, cyan) and overlay of all channels (scale bar is 5μm), **c**, Quantitative representation of changes in cytoskeletal (total tubulin volume, μm^3^ and surface area, μm^2^) and mitochondrial (total volume, μm^3^) features, as well as organelle polarity during adaptation to culture (significant differences shown in graph), **d**, Heatmap showing changes in abundance of markers of HSC quiescence (H1-0), activation and proliferation (CDK6, MKI67, CCNB1, AURKB), mitochondrial proteins (TFAM,TIMM23,TOMM22), structural cytoskeletal proteins (alpha-tubulin:TUBA1B, TUBA1C) and regulators of cell polarity (CDC42, SEPTIN7, DOCK7/11) in HSCs grown in basal media across days in culture from total proteomics, showing similar trends as microscopy analysis.

In agreement with proteomic profiles, progressive restructuring of intracellular architecture could be observed across culture time points and during transition from quiescence to a proliferative state (Figure 4b, c). Consistent with the proteomic evidence for cytoskeletal remodeling, total tubulin volume and surface area significantly increased over time, indicating coordinated remodeling and expansion of the microtubule scaffold during early adaptation and later expansion stages (Figure 4c, d). Day 3 cells displayed an intermediate architectural state, coinciding with the onset of proliferative activation (Figure 1b), as supported by the loss of the quiescence marker H1-0 and increase in regulators of active cell cycle and proliferation including CDK6, MKI67, CCNB1 and AURKB (Figure 4d). Such architecture may reflect a discrete phase of cortical redistribution or cytoskeletal planarization, consistent with structural reorganization accompanying entry into cell cycle^36,37^. In parallel with the cytoskeletal remodeling, mitochondrial organization shifted from a fragmented and spatially polarized distribution at day 0 toward a more elongated and globally redistributed network by day 7 (Figure 4b). This structural change coincided with a significant increase in total mitochondrial volume (Figure 4c) and mitochondrial markers in proteomic profiles (TFAM, TIMM23, TOMM22) (Figure 4d), suggesting a coordinated expansion and remodeling of organelle architecture alongside microtubule and cytoskeletal structures.

To determine how cytoskeletal and organelle changes affected the spatial organization within the cell, intracellular polarity was quantified based on 3D volumetric analysis of organelle localization, focusing on mitochondria as organelles that are actively re-positioned by microtubule-motor systems and have been associated with HSC activation states^38–40^. This analysis revealed a progressive reduction in polarity across culture timepoints, with D7 cells exhibiting significantly lower polarity (Figure 4c). A heterogeneous polarity phenotype was observed among D3 cells, with a subset retaining asymmetric organization while others showed reduced polarity, consistent with a transitional cell cycle state. Interestingly, and in line with the imaging-based polarity measurements, proteomic profiling revealed coordinated temporal changes including an early enrichment followed by gradual loss of CDC42 and SEPTIN7 (Figure 4d), which form a central axis in governing cell polarity and stem cell function in HSCs^23,24,41^. This shift was concurrent with a gradual loss of DOCK7 and increase in DOCK11(Figure 4d), two regulators of RAC/CDC42 activation states, suggesting a switch in small GTPases that control the polarity signaling network^23,24,42,43^. While the CDC42/Septin7 axis supports HSC polarity and function^24^, the late-stage upregulation of DOCK11 can facilitate a global, symmetric activation of CDC42 that mirrors the aged HSC phenotype^42^, where the loss of spatial signaling leads to volumetric expansion and the permanent dismantling of the polarized stem cell architecture.

Together, these spatial and proteomic data reveal the underlying cytoskeletal remodeling during HSC adaptation and expansion that ultimately results in the hallmark of *ex vivo* HSC expansion; cell division coupled with loss of polarity and stemness or emergence of aging-associated signatures.

### Cytoskeletal and mitochondrial morphodynamics define polarity trajectories during adaptation

To capture the global architectural remodeling that occurs during *ex vivo* culture, we performed integrated morphological profiling based on tubulin and mitochondrial features but excluding polarity. Dimensionality reduction demonstrated that variation across cells was dominated by coordinated changes in cytoskeletal and mitochondrial features, including total volume, surface area, and branching complexity (Figure 5a), indicating that structural scaling of these compartments represents a principal axis of HSC adaptation and expansion in culture.

**Figure 5.**
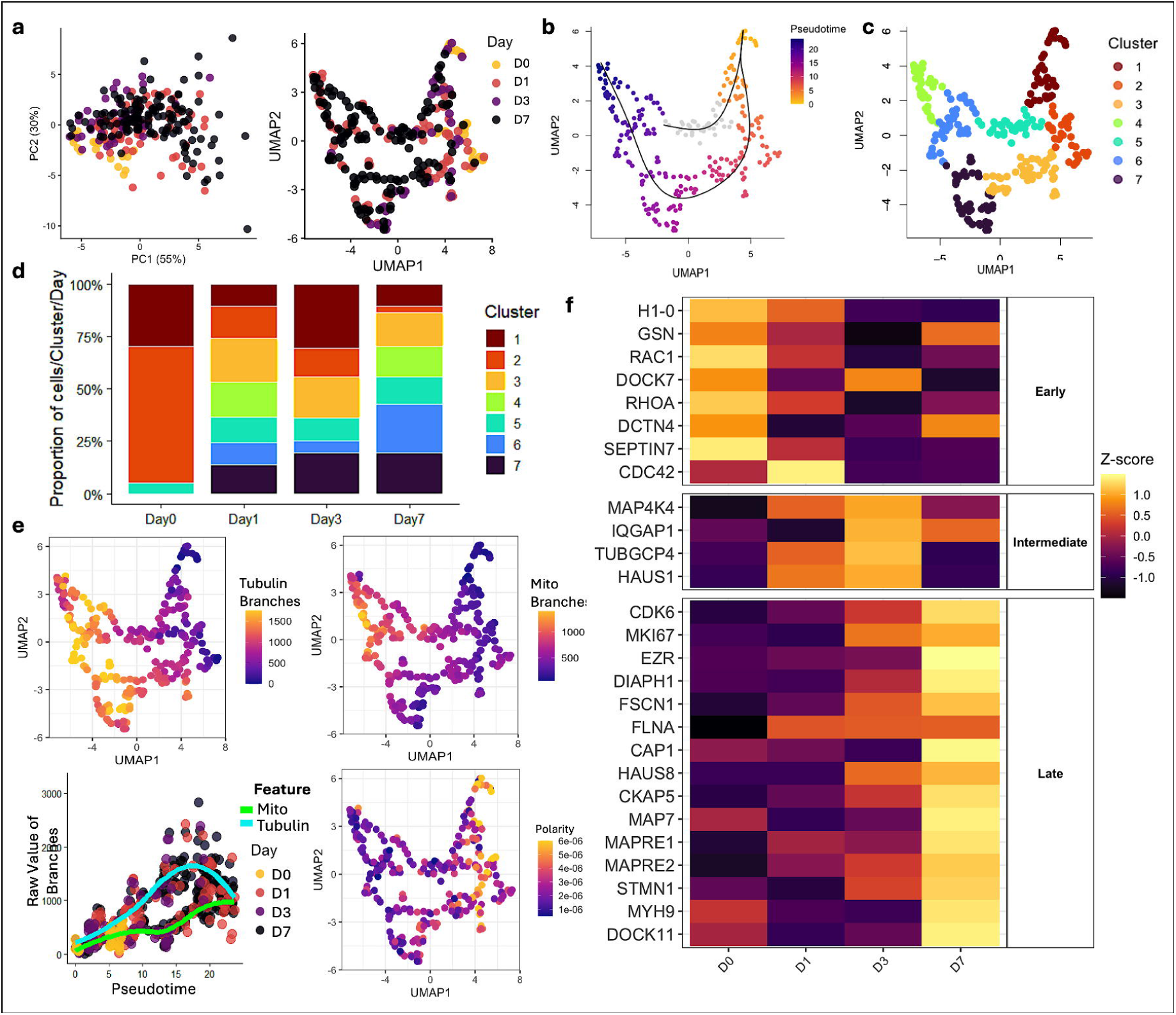
Cytoskeletal and mitochondrial morphodynamics during HSPC adaptation. **a,** PCA and UMAP dimensionality reduction based on morphological features of HSPCs showing unique day-specific patterns, **b, c**, Pseudotime and unsupervised clusters generated based on morphological features, **d**, Proportion of cells from each morphological cluster comprising total cell population on each day, **e**, Overlay of morphological features on the UMAP indicating an association between morphological features and pseudotime, **f**, Scaled expression of proteins associated with regulation of polarity, microtubule branching, cytoskeletal maturation, and membrane and organelle interaction from total proteomics of HSCs across days in culture.

Unsupervised clustering combined with trajectory inference mapped a continuous pseudotime from quiescent HSCs (predominantly day 0, Clusters 1 and 2) to late-stage cultured cells mainly from day 7, Cluster 4, consistent with a gradual transition from a primitive to a structurally mature cellular state (Figure 5b-d; Supplementary Figure 6a-c). Along this trajectory, cytoskeletal branching gradually increased and preceded mitochondrial branching and network complexity (Figure 5e), supporting a model where early structural remodeling underlies the metabolic transitions traditionally associated with rising energetic demand, metabolic maturation or loss of function in HSCs^44–48^. Primitive cells at the start of the trajectory exhibited fragmented mitochondria and minimal cytoskeletal complexity, consistent with a low-energy quiescent state^40^, whereas later-stage cells displayed expanded cytoskeletal architecture together with elongated and increasingly interconnected mitochondrial networks, in line with a high-energy demand mature state^46^ (Figure 5b-c; Supplementary Figure 6b-c). Notably, cytoskeletal branching increased (cluster 7) prior to expansion of mitochondrial branching (clusters 6), indicating that remodeling of the cytoskeleton precedes mitochondrial network scaling (Figure 5e). Maturation of cytoskeletal architecture coincided with increase in mitochondrial branching complexity at later stages of culture progression (Figure 5e; Supplementary Figure 7a). An alternative trajectory showed moderate levels of cytoskeletal complexity and mixed mitochondrial morphology, consistent with either cycling cells or other populations that did not fully progress along the dominant adaptation and expansion path (Figure 5b, c). Interestingly, overlaying polarity on the pseudotime map indicated a progressive decline in polarity along the structural trajectory (Figure 5e; Supplementary Figure 7b), linking mitochondrial morphology and distribution directly to the loss of intracellular asymmetry observed during culture adaptation and expansion. This is in line with previous work reporting that mitochondrial morphology and localization can reflect quiescence or activation of mouse HSCs in the bone marrow^38^, further supporting a tight link between cytoskeletal and mitochondrial remodeling, polarity reorganization and maintenance of quiescence or stemness in HSCs.

To resolve the molecular mechanism underlying this trajectory, we analyzed the expression of regulatory and structural proteins in HSCs during adaptation and expansion in culture (Figure 5f). The early phase (D0/D1) was defined by an enrichment of proteins that maintain flexible (GSN), asymmetric cytoplasmic organization (RAC1/CDC42/SEPTIN7 axis). The intermediate stage (D3) was characterized by an increase in regulators of scaffold-integration (IQGAP1), supporting a transition toward cytoskeletal branching (MAP4K4, UBGCP4, HAUS1). In contrast, the late timepoint (D7) showed a coordinated shift toward membrane-anchored structural maturation (EZR, DIAPH1). This was accompanied by higher recruitment of actin-crosslinking and bundling components (FLNA, FSCN1, MYH9) and microtubule-associated proteins (CKAP5, MAP7, STMN1). Analysis of cytoskeletal and mitochondrial structural components further supported this remodeling trajectory (Supplementary Figure 7c). Early adaptation was defined by polarity-supporting cortical scaffold elements and intermediate filament (spectrins and vimentin), coupled with a reservoir of fragmented mitochondria (TFAM, VDAC1, and FIS1) that are positioned towards distal regions, potentially via RHOT1/MIRO1^49,50^, in line with polarized high mitochondria mass quiescent HSCs^39,40^. As cells transitioned to the intermediate stage, they showed a synchronized increase in actin-branching machinery (ARPC2/3, TUBGCP4) and mitochondrial complexes (TOMM22, OXA1L), signaling the start of a coordinated structural and metabolic reorganization. This maturation culminated at day 7 with the progressive enrichment of stabilized microtubule and contractile actomyosin components (TUBB3, ACTN2/4, VCL), as well as high OXPHOS machinery (IMMT, TIMM23, ATP5PF) where active symmetric mitochondrial distribution may be facilitated by RHOT2 (MIRO2), as observed in our 3D imaging. Collectively, these temporal shifts further support that remodeling of the cellular scaffold precedes and facilitates the expansion and maturation of the mitochondrial network during *ex vivo* HSC adaptation and expansion.

### FASN inhibition uncouples cytoskeletal and polarity trajectories during culture adaptation

To decipher the significance of early FASN increase and its role in cytoskeletal and organelle remodeling, we targeted the initial window of HSC adaptation using Denifanstat (TVB-2640), a phase II clinical FASN inhibitor. We found that early modulation of FASN by low dose of the inhibitor, strictly during the adaptation phase (day 0), significantly increased the number of CD34^+^ progenitors and HSCs by day 7 (Figure 6a; Supplementary Figure 8a, b). In contrast, initiating FASN inhibition on day 3, when the cells have passed the early adaptation window and progressed into the cell cycle towards a proliferative state, yielded no significant effect (Supplementary Figure 8c). This is in line with our integrated proteomic and imaging analyses, verifying a narrow temporal window where FASN activity is required to govern early adaptation, independent of its role in supporting later expansion.

**Figure 6.**
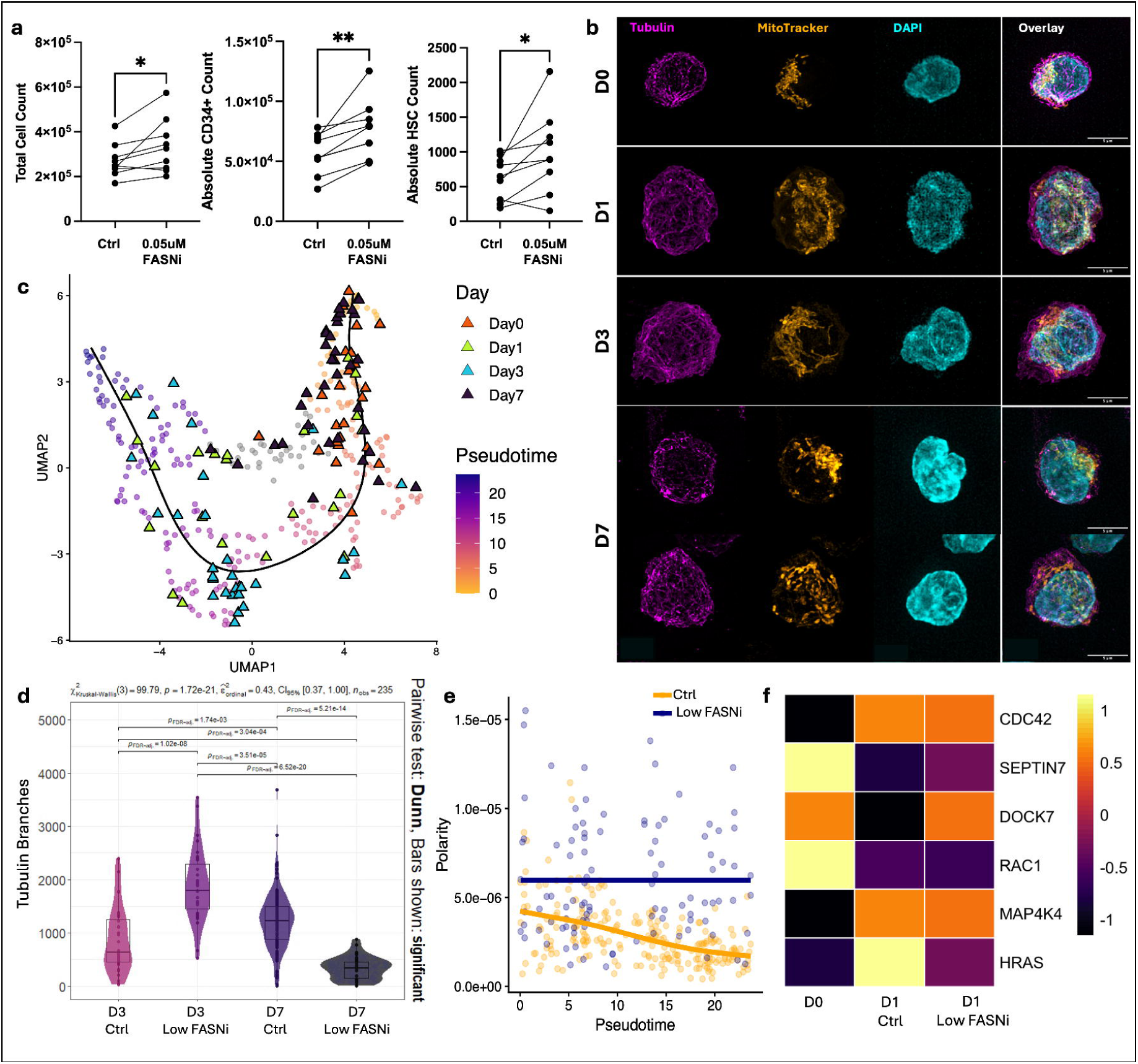
FASN inhibition with TVB-2640 disrupts cytoskeletal and mitochondrial remodeling and alters morphological trajectories during culture adaptation. **a,** Changes in cell count across total cells, CD34^+^ cells and phenotypic HSCs from control untreated vs FASN-inhibited cells treated with TVB-2640 initiated on day 0 (one-way ANOVA was used to calculate significance, * = p<0.05, ** = p<0.005), **b**, Representative microscopy images of FASNi cultures demonstrating changes in tubulin (magenta), mitochondrial features and polarity (MitoTracker Red, orange), nucleus (DAPI, cyan) and overlay of all channels across days in culture (scale bar is 5μm), **c**, Overlay of FASNi cells on reference pseudotime trajectory generated based on morphological features of untreated cells, **d**, Comparative analysis of tubulin branching in D3 and D7 FASNi vs untreated HSCs (significant differences and p values (p.adj <0.05) shown in graph) **e**, Polarity trajectories over pseudotime for untreated and FASNi cultures with GAM-smoothed trends shown for each group, **f**, Proteomic changes associated with early polarity and cytoskeletal remodeling in HSCs in D0 vs D1 untreated or FASNi cells.

To determine whether inhibition of FASN translated into morphological changes, we performed super-resolution imaging following FASN inhibition and quantified cytoskeletal and mitochondrial features (Figure 6b; Supplementary Figure 8d). Treated cells were projected onto UMAP and pseudotime models trained on morphological features from untreated control cells (Figure 6c). Day 0 and day 1 FASNi-treated cells overlapped extensively with their untreated counterparts (Figure 6c). By contrast, a larger proportion of day 3 FASNi-treated cells localized to later pseudotime positions, whereas many day 7 cells shifted back toward earlier pseudotime, co-clustering with day 0 cells in the UMAP embedding, indicating a more primitive morphology with low cytoskeletal complexity and fragmented mitochondrial (Figure 6c, Supp. Fig. 8d). This redistribution across the pseudotime predominantly drove FASNi-treated cells to later pseudotime at day 3, mapping to Clusters 6 and 7 (Figure 6c), indicating a complex branched cytoskeletal network (Figure 6d). Despite the increased cytoskeletal complexity, the cells maintain a moderate level of polarity and lose the morphological maturity associated with complex cytoskeletal-mitochondrial network (cluster 4) (Figure 6e; Supplementary Figure 9a-d), suggesting a disconnect between cytoskeletal and organelle remodeling under FASNi treatment. The structural changes downstream of FASN inhibition corresponded with an uncoupling of the pattern of polarity decline observed in the untreated cells, resulting in improved maintenance of polarity among day 7 FASNi-treated cells (Figure 6b, e). Indeed, FASN inhibition reinforced polarity at day 7 (Figure 6b, e; Supplementary Figure 9a, b), consistent with our earlier observations of FASN-dependent regulation of cytoskeletal and organelle dynamics (Figure 3d). Overall, loss of early FASN-mediated cytoskeletal remodeling compromises the scaffolding and microtubule anchoring required for extensive mitochondrial expansion or distribution, resulting in retention of polarity, ultimately altering cell division and expansion trajectory (Supplementary Figure 9e).

To resolve the underlying molecular mechanisms, we applied our low-input proteomics analysis to HSCs during this critical early transition window (Supplementary Figure 9f, g). Despite the overnight increase in mTORC1 signaling and E2F targets regardless of treatment, FASN inhibition led to a significant change in KRAS_DN GSEA signature (Supplementary Figure 9h). Interestingly, changes in CDC42, SEPTIN7, RAC1 and MAP4K4 followed the established early adaptation patterns, independent of treatment (Figure 6f). However, DOCK7 and HRAS in D1 FASNi-treated cells showed levels similar to D0 cells (Figure 6f), in line with altered regulation of RHO effector signaling, cytoskeletal remodeling and polarity. In combination with the expansion outcome from early versus late FASN inhibition, these data reveal that FASN acts as an early metabolic rheostat during the adaptation phase, specifically orchestrating cytoskeletal-mitochondrial interactions and modulating RAS signaling to dictate the trajectory of HSC *ex vivo* expansion, entirely independent of cell division. Collectively, these findings position the FASN-RAS axis as a dynamic coordinator linking early metabolic adaptation to structural transitions required for proliferation, defining a signaling framework in which cellular division could proceed without complete loss of polarity. Importantly, and in line with previous studies emphasizing the role of cell-cycle independent adaptation in *ex vivo* HSC culture^5^, our data places polarity remodeling downstream of metabolically regulated cytoskeletal changes, rather than an inevitable consequence of cell-cycle entry, identifying the FASN–RAS axis as an early regulator of architecture polarization during cytokine-driven HSC expansion.

### Pharmacological perturbation of tubulin dynamics recapitulates the downstream cytoskeletal control of HSC adaptation

To independently validate the role of cytoskeletal adaptation, we directly targeted microtubule polymerization using Vincristine, a clinical inhibitor of tubulin assembly that acts independent of FASN-mediated cytoskeletal modifications. Low concentrations of Vincristine enhanced expansion of phenotypic HSCs, whereas medium to high doses suppressed both total proliferation and maintenance of HSCs (Figure 7a). The morphological changes during culture were assessed using our super-resolution microscopy analysis (Figure 7b). At early time points, day 1 low Vincristine-treated cells closely overlapped with untreated controls in pseudotime projections and showed no significant alterations in tubulin or mitochondrial volume (μm^3^) and branching (Figure 7c, Supplementary Figure 10a). By contrast, day 7 low Vincristine-treated cells were more restrained, shifted toward earlier pseudotime positions, exhibited a reduction in tubulin and mitochondrial complexity while maintaining polarity, overall showing less mature phenotypes as evident in minimal representation in Cluster 4 (Figure 7c-e; Supplementary Figure 10a-c). However, the medium or high doses of Vincristine did not promote cell growth (Figure 7a) and showed accumulation of disorganized tubulin and altered mitochondrial morphology (Supplementary Figure 10d), consistent with cytoskeletal collapse and cell-cycle arrest. Together, our data establishes cytoskeletal remodeling, not only as a structural requirement for mitochondrial expansion, but also as a regulatory axis during the adaptation phase that directs the *ex vivo* HSC expansion trajectory.

**Figure 7.**
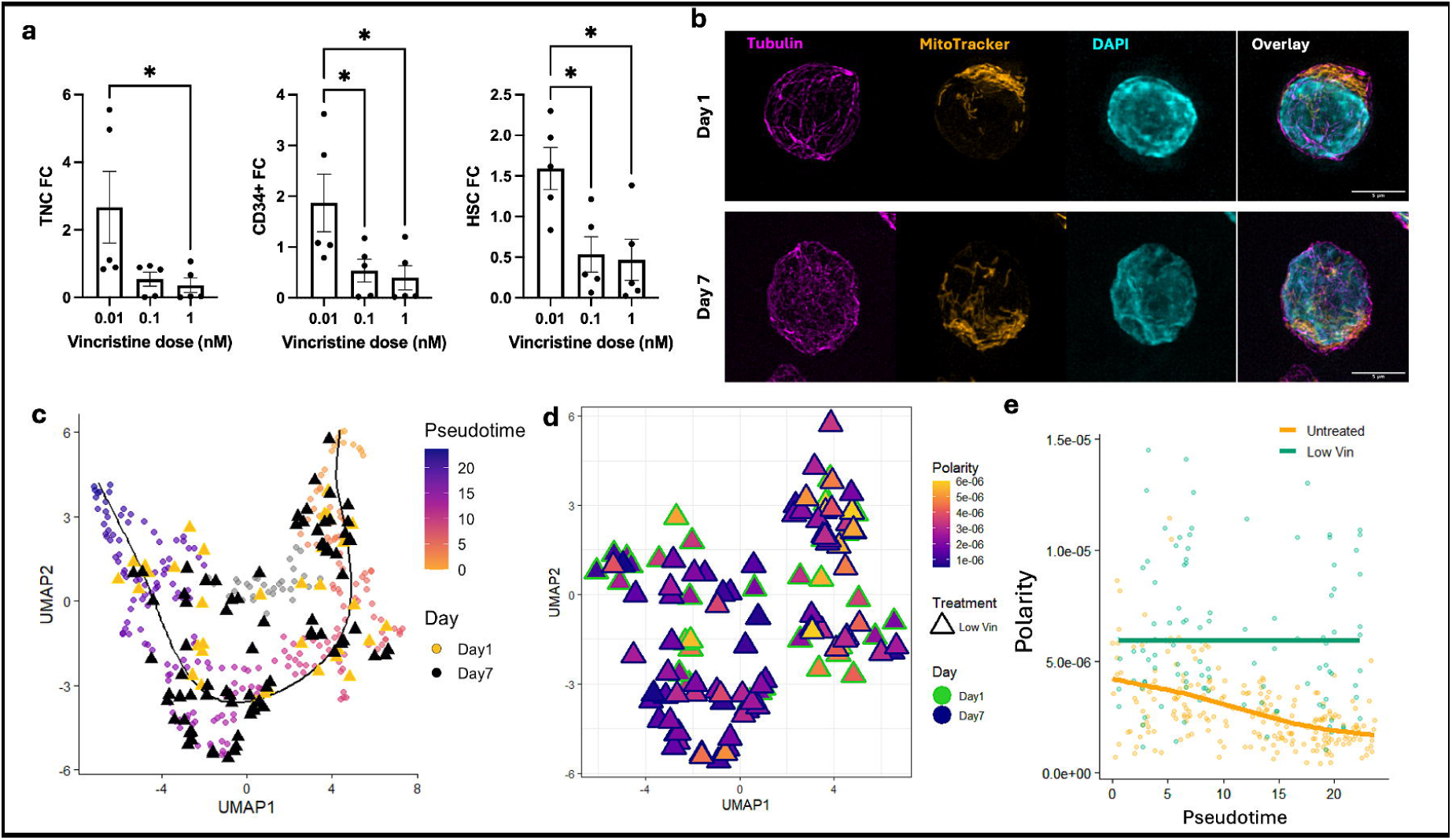
Inhibition of cytoskeletal remodeling alters HSPC adaptation trajectories. **a**, Fold change (FC) of total nucleated cells (TNC), CD34^+^ cells and phenotypic HSCs (vs untreated control) following treatment with low (0.01nM), medium (0.1nM) or high (1nM) dose of Vincristine (one-way ANOVA was used to calculate significance, * = p<0.05) **b**, Representative microscopy images of cells treated with low dose of Vincristine indicating morphological changes in tubulin (magenta), mitochondrial features and polarity (MitoTracker Red, orange), nucleus (DAPI, cyan) and overlay of all channels across days in culture (scale bar is 5μm), **c,** Overlay of Vincristine-treated cells on reference pseudotime trajectory from untreated cells, **d,** Overlay of polarity for the Vincristine-treated cells projected over reference pseudotime, labelled by treatment and days in culture, **e,** Polarity trajectories over pseudotime for untreated and Vincristine-treated cultures with GAM-smoothed trends shown for each group.

## Discussion

By integrating ultra-low input quantitative proteomics with super-resolution imaging, we define a metabolic-cytoskeletal axis that governs HSC adaptation and directs the trajectory of *ex vivo* expansion. This proteomic signature is established within the first 24 hours, independent of cell division, in line with other reports indicating that the loss of polarity and stemness is an early adaptive outcome rather than a byproduct of proliferation^5^.

While the role of RAS and RHO GTPases in regulating HSC polarity, cell cycle and fate is well-documented in murine models^18^ and during cell division^20^, their function in human HSCs during early culture adaptation has remained elusive. Our approach reveals how biosynthetic flux, cytoskeletal architecture, and organelle organization are functionally coupled to support cell behavior and fate during the adaptation process. At the proteomic level, we show a metabolic switch, particularly lipid biosynthesis via FASN, coincides with changes in RHO-family effectors (RAS, RHOA, RAC1) and cytoskeletal remodeling as well as a loss of polarity via the SEPTIN7/CDC42 axis. Our imaging corroborates this, showing that progressive cytoskeletal and mitochondrial branching complexity mirrors the metabolic reprogramming and loss of polarity captured in our proteomics and reported previously^38,46^. Together, these data establish a metabolic-cytoskeletal continuum that initiates during early adaptation to determine the eventual HSC *ex vivo* expansion trajectory.

This metabolic-cytoskeletal coupling suggests that lipid synthesis extends beyond providing metabolic and membrane substrates^51,52^ acting as a regulator of subcellular organization by modulating signaling and cytoskeletal proteins^35,53^. Sustained abundance of FASN throughout culture, along with identification of a narrow temporal window when FASN inhibition can divert the morphological and *ex vivo* expansion trajectories, confirm that the principal regulatory shift in this process is not simply lipid substrate availability but an early programmed rewiring of the signaling and cytoskeletal network. The coupling of *de novo* fatty acid synthesis with microtubule and cytoskeletal organization, consistent with the role of palmitoylation in stabilizing tubulin and RHO-family effectors^35,54^, positions lipid metabolism as an early metabolic rheostat that calibrates the overall physical state of the cell during adaptation to culture.

Pharmacologic inhibition of FASN disrupts this coordinated metabolic-cytoskeletal remodeling, revealing a reversible checkpoint during adaptation, linking lipid biosynthesis to structural remodeling and expansion outcomes. Restriction of tubulin polymerization by Vincristine provides an orthogonal validation of the role of cytoskeletal remodeling in directing HSC expansion trajectories during adaptation. However, excessive perturbation of microtubules results in structural disintegration. Overall, our data demonstrates the central role of cytoskeletal dynamics in maintaining the integrity of cellular architecture, polarity and cell competence to adapt to *ex vivo* culture, underscoring the significance of finely tuned cytoskeletal changes in sustaining HSC polarity and self-renewal potential.

Collectively, these findings establish a metabolic-cytoskeletal axis that determines cellular polarity or a/symmetry prior to the onset of cell division. This suggests that polarity loss during cytokine-driven culture is a regulated feature of early cellular adaptation, rather than an inevitable consequence of cell-cycle entry. Our study offers a “biology-first” roadmap for developing new approaches to preserve primitive HSC identity during *ex vivo* expansion, thereby enhancing the effectiveness of HSCs as a curative platform for cellular and gene therapies.

## Methods

### KG1a cell culture

The KG1a cell line was obtained from ATCC. Cells were quickly thawed in a 37°C water bath and washed once with PBS + 10% fetal bovine serum (FBS). Cells were cultured in RPMI supplemented with 20% FBS, 100 units/mL Penicillin-Streptomycin and 2mM L-glutamine and maintained at 1-1.5 x 10^6^/mL of media in a humidified incubator at 37°C with 5% CO_2_. Cells were confirmed as Mycoplasma free by monthly PCR checks and were tested for expression of CD34 by flow cytometry at the start of the culture and at random intervals. Cells were washed with PBS, centrifuged at 400 g for 5 minutes and resuspended in PBS + 1% FBS prior to sorting for proteomics analysis.

### Umbilical Cord Blood preparation

Umbilical Cord Blood units (UCB) were obtained from Anthony Nolan with full informed consent under HTA license (22527). Ethical approval for use of human tissue for research was granted by the NRES committee (NHS research ethics service - East Midlands-Derby ref No. 20/EM/0028) and the London Bridge Ethics Committee (ref No. 19/LO/1411). Mononuclear cells were isolated by density centrifugation using Ficoll-Paque (Cytiva). CD34^⁺^ cells were positively selected using Miltenyi anti-CD34 conjugated magnetic beads and MACS columns following manufacturer’s instructions (Miltenyi Biotech). After washing cells were resuspended in Cryostor CS10 (StemCell Technologies), aliquoted and cryopreserved. On the day of experiments, aliquots were rapidly thawed in a 37 °C water bath and washed twice with PBS + 2% FBS (hereafter referred to as FACS buffer), before counting and pooling. Each pool contained a minimum of 3 individual samples. Pools were seeded in appropriate culture wells (see below) for experiments and remaining cells were stained with HSPC phenotyping antibodies (see key resources table) for sorting and proteomics.

### Primary CD34^+^ cell culture conditions

All CD34^+^ cell cultures were carried out in basal media containing StemSpan SFEM II (StemCell Technologies) supplemented with recombinant human stem cell factor (SCF; 150ng/ml), recombinant human FLT3 ligand (FLT3L; 150ng/ml) and recombinant human thrombopoietin (TPO; 20ng/ml) (all from Peprotech). Cells were incubated in a humidified incubator at 37 °C with 5% CO_2_, with a media change on day 3, for up to 7 days. In conditions where UM729 is indicated, this was added as a supplement to basal media at 0.5 µM concentration. To inhibit FASN (FASNi), media was supplemented with Denifanstat (TVB-2640) at 0.05 or 5µM concentration at day 0 and re-supplemented with media change on day 3 or only added on day 3. To inhibit tubulin polymerization, Vincristine was added to the media at concentrations of either 1nM, 0.1nM or 0.01nM on day 0 of culture and re-supplemented along with media change on day 3.

### Flow cytometry staining, analysis and sorting

Cells were stained for 30 minutes at 4 °C in FACS buffer containing the indicated antibodies to phenotype HSCs as: day 0 and day 1 in culture HSC^1^ (CD34^+^ CD38^low^ CD45RA^-^ CD49f^+^ EPCR^+^) or HSC^2^ (CD34^+^ CD38^low^ CD45RA^-^ CD49f^+^ EPCR^-^), and day 3 and 7 in culture HSC^e^ (CD34^+^ CD45RA^-^ CD90^+^ EPCR^+^), adjusting the panels to account for well-documented surface marker alterations in immunophenotypic *de novo* cord blood and *ex vivo* expanded HSCs^58–62^ (antibody information detailed in the resources table). Following wash and centrifugation in FACS buffer (400 g, 10 minutes), the cells were resuspended in FACS buffer containing Sytox green as cell permeable live dead dye, and data was acquired on CytoFlex flow cytometers and sorters (LX375 or SRT sorter, Beckman Coulter). For all flow cytometry analyses, cells were initially identified based on forward and side scatter. Dead cells were excluded based on Sytox green staining. Full gating strategies for each time point are reported in Supplementary Figure 2.

### Low-input proteomics sample preparation

For efficient low-input proteomics, 100 µL PCR tubes were filled with 0.2% n-dodecyl-β-D-maltose (DDM) and left to coat overnight at 4 °C. Lysis buffer was prepared fresh at 2X concentration, containing 0.4% DDM, 2 mM sodium orthovanadate, 2 mM beta-glycerophosphate, 80 mM tetraethylammonium bromide (TEAB, pH= 8.5). Tris (2-carboxyethyl) phosphine hydrochloride (TCEP) was dissolved fresh and added to the lysis buffer at 12 mM prior to use. All tubes and sample handling steps were confined to a laminar flow hood, excluding cell sorting, and were arranged to minimize light exposure. Coating buffer was completely removed prior to addition of 2 µL of the lysis buffer for either 50 or 200 cells or 5 µL for 2000 cells, respectively. Cells were sorted directly into the lysis buffer using a CytoFLEX SRT Cell Sorter. Samples were quickly spun down and stored on dry ice and transferred to -80 °C until further processing.

For proteomics preparation, samples were removed from -80 °C, quickly spun down and lysis was performed at 70 °C for 30 minutes with a heated lid at 105 °C using a thermocycler. When isotonic sheath fluid was used during cell sorting, 1μL of water was added to the samples prior to lysis. However, this step was not required when the sheath fluid was hypotonic. Samples were cooled down to undergo alkylation using iodoacetamide (IAM) at final concentration of 100 mM at room temperature (RT) in the dark for 15 minutes. To prepare the enzymatic digestion cocktail, 100 ng/mL Trypsin-LysC dissolved in acetic acid-based buffer was diluted 1:10 in 200 mM TEAB containing 25 units/μL ultrapure benzonase nuclease. Either 1µl of the enzymatic cocktail was used for 50 or 200 cells or 4 µL of the cocktail was added to 2000 cells prior to initiating a 3-hour digestion at 37 °C using a thermocycler with a heated lid at 105 °C. The pH of the samples was adjusted using 10% Trifluoroacetic. Urea lysis buffer was prepared at a final concentration of 9 M urea, 1 mM sodium orthovanadate, 1 mM beta-glycerophosphate, 2.5 mM sodium pyrophosphate and 20 mM 4-(2-hydroxyethyl)-1-piperazineethanesulfonic acid (HEPES, pH= 8.0). Lysis buffer addition, cell sorting and sample storage procedures followed the DDM-lysis process; however, lysis was performed using cooled water bath sonication for 10 minutes. TCEP and IAM were used, at the same concentration as DDM-lysis, respectively for reduction at 55 °C for 15 minutes followed by alkylation at RT in the dark for 15 minutes. Samples were then diluted with 20 mM ammonium bicarbonate to reach a urea concentration of 2 M prior to enzymatic digestion as described earlier, followed by peptide acidification and volume adjustments using 10% or 0.1% TFA, respectively. LC-MS water was used throughout all the processes involving sample preparation for proteomics.

Acidified peptides were loaded onto EvoTip Pure tips for nanoUPLC using an EvoSep One system and following the manufacturer’s protocol. A pre-set 60SPD gradient was used with an 8 cm EvoSep C_18_ Performance column (8 cm x 150 mm x 1.5 mm).

The nanoUPLC system was interfaced to a timsTOF HT mass spectrometer (Bruker) with a CaptiveSpray ionisation source. Positive PASEF-DIA, nanoESI-MS and MS^2^ spectra were acquired using Compass HyStar software (version 6.2, Bruker). Instrument source settings were capillary voltage, 1,600-1,700 V; dry gas, 3 l/min; dry temperature; 180 °C. Spectra were acquired between m/z 100-1,700. DIA windows were set to 25 Th width between *m/z* 400-1201 and a TIMS range of 1/K0 0.6-1.60 V.s/cm^2^. Collision energy was interpolated between 20 eV at 0.65 V.s/cm^2^ to 59 eV at 1.6 V.s/cm^2^.

### Protein search and identification

LC-MS data, in Bruker.d format, was processed using Spectronaut (version 19 - Fig. 1 data or 20 - Fig. 3 data) software and searched in direct-DIA mode against the human subset of SwissProt appended with common proteomic contaminants. Search criteria specified: enzymes, trypsin/P; variable modifications; acetyl (protein N-term), oxidation (M); fixed modification, carbamidomethyl; digest type, specific; max peptide length, 52; min peptide length, 7; missed cleavages, 2; MZ extraction strategy, maximum intensity; precursor q-value cutoff, 0.01. DIA XIC extraction windows for IM, IT, MS^1^ and MS^2^ were all set as dynamic. Identifications were filtered to require protein q-values < 0.01 and a minimum of two peptides per accepted protein. Normalized, DIA-derived MS intensities were exported from Spectronaut for differential abundance testing.

### Proteomics data analysis and statistics

Calculation of log2 fold change and differential abundance testing was performed using limma via FragPipe-Analyst^63^. Proteins analyzed were filtered by presence in at least 75% of samples in at least one group. Zero imputation was applied and adjusted p-values were calculated using Benjamini-Hochberg correction for temporal analysis or local and tail area for FASN inhibition analysis. Subsequent biological analyses are further described in dimensionality reduction-based analysis, presented through an online portal (https://shiny.york.ac.uk/ProteoStem_Viewer/) and coding pipelines are provided on Zenodo (https://zenodo.org/records/20204118).

### Super-resolution microscopy sample preparation and image acquisition

Cells from pools of five to fifteen individual UCB units were seeded into ibidi-8-well-chamber-slides and incubated for either 1 hour on day 0 or cultured and treated as described earlier only using the basal media without UM729. Cells were incubated with MitoTracker™ Red CMXRos at final concentration of 200 nM for 30 minutes, to stain all mitochondria regardless of oxidative activity. Cells were fixed by adding equal volume 8% methanol-free formaldehyde in 2x cytoskeletal buffer (CB) containing 20 mM MES (pH=6.1), potassium chloride (280 mM), magnesium chloride (6 mM), and EGTA (34 mM) for 20 minutes at 37 °C. Permeabilization and autofluorescence quenching were achieved by incubation in 0.5% Triton 100X for 5 minutes and 0.1 M glycine for 10 minutes, in CB at RT. Samples were washed twice in TBS containing 0.5% Tween 20 (TBST) and blocked for 1 hour at RT using bovine serum albumin (BSA) final concentration of 4% in TBST. Following overnight incubation with a primary alpha-tubulin antibody in the 2% BSA in TBST at 4 °C, samples were washed 3 times with TBST and incubated for 2 hours at RT with a secondary antibody in blocking buffer containing DAPI for nuclear staining. Following 3 washes, the samples were covered with anti-fade gold, left to cure overnight and stored at 4 °C prior to imaging by lattice-structured illumination microscopy (SIM) on a Zeiss Elyra 7 platform.

### Image analysis pipelines

#### A) SIM^2^ processing

All images were SIM^2^ processed in ZEN Black microscopy software with processing strength default presets applied based on the fluorophore signal intensities of the central Z slice and the processed output scaled to the original raw intensity values. Weak-fixed, standard-fixed, and low-contrast processing presets were used for MitoTracker Red (signal intensities 800-1000 grey levels), CoraLite^®^ Plus 647 (signal intensities above 1500 grey levels) and DAPI, respectively. All image acquisition and processing settings were maintained constant across all experiments.

#### B) .czi cleanup and pre-processing steps

SIM^2^-processed images were opened in Fiji, and the XY centers of cells were identified based on the nuclear channel signal, following a series of median and Gaussian filtering processes and Otsu-based thresholding. A size filter was set to exclude debris and validate cell centers. These centers were used as the midpoint of a 512 px x 512 px selection box to crop and generate individual .tiff Z-stacks from the original .czi file. In each Z-stack .tiff, cell segmentation was achieved by signal thresholding, by empirically assigning a signal to noise ratio threshold of 3, followed by using a series of morphological filters, and determining the first and the last slices along the Z axis, allowing for additional 5 slides in each direction. Images containing at least 45 Z-slices were considered a valid stack.

#### C) Mitochondrial and Tubulin Analysis

Validated trimmed 16-bit Z-stacks .tiff files were converted to 8-bit images, without contrast modification, and were analyzed using 3D thresholding in the Mitochondria Analyzer plugin in Fiji to generate masks for tubulin and mitochondria. Block and c-value were selected for optimal mitochondrial and tubulin detection and were maintained constant across experiments. Masks were QC’d and used to run 3D Analysis batch function to obtain “per-cell” statistics. The data is available in Supplementary data file Table 1.

#### D) Polarity assessment

An isotropic image was generated from each validated trimmed Z-stack by setting uniform voxel widths, heights, depths and linear interpolation of the Z slices in the stack. Binary masks were generated for cells using the MorphoLibJ plugin in Fiji on the max projection from all channels. This region was extended to the entire 3D Z-stack to define the cellular area and a value of zero was assigned to the background defined as the field outside this cellular area. Polarity was evaluated following the dipole-based algorithm presented by Schuster et al.^64^ generated by P = q_+ve,norm_d_norm_, where q_+ve,norm_ is the normalized positive fluorescent charge of the cell and d_norm_ is the distance between the positive and negative barycenters, normalized by the maximum diameter of the XY plane of the cell.

In summary, the average fluorescent intensity of the 3D cellular area was normalized to 1 and was subtracted from the actual intensity value of each pixel to create positive and negative regions in the cell. The centers of gravity for the positive and negative regions were determined using charge- and bit-based, as well as a distance-based normalization based on the maximal XY radius of the cellular mask. This was used to calculate cell size-normalized dipole moments, which was used as a measure of polarity.

### Dimensionality reduction-based analysis

The most abundant variations in temporal changes in proteomics or morphological features, excluding polarity, were investigated using principal component analysis (PCA). More complex relationships between various populations were explored using uniform manifold approximation and projections (UMAP) via the uwot package. The PCA and UMAP projections were used to perform unsupervised clustering and infer pseudotime using the Cluster and Slingshot packages.

Proteomics analysis was followed by investigating the statistically significant temporal changes in protein abundance along the pseudotime via Tradeseq. Additionally, differentially abundant proteins were clustered based on their real-time changes in pattern of expression using the DEGreport package. Functional relevance of all these differentially abundant proteins was further evaluated by gene ontology overrepresentation or gene set enrichment analysis (GSEA) using the ClusterProfiler package.

For morphological features, the resulting unsupervised clusters within the untreated samples were assigned back to the samples to enable unbiased statistical analysis of changes in tubulin, mitochondrial and polarity features between the clusters and across the pseudotime to infer functional and biological relevance. Furthermore, the UMAP model trained based on the untreated samples was applied as the reference to generate UMAP and determine pseudotime for Denifanstat- or vincristine-treated samples.

Nonparametric comparisons across morphological clusters were performed using Kruskal Wallis test followed by Dunn’s nonparametric pairwise comparison, with p-value and adjusted false discovery rate (FDR) <0.05 considered significant, respectively. The same statistical analyses and significance threshold were applied to compare each selected morphological feature between untreated and treated samples. p-values and FDRs are reported within the graphs for each comparison as generated by the ggstatsplot package.

### Data availability

Proteomics data has been deposited at MassIVE (MSV000099647) and referenced in ProteomeXchange (PXD069943), made available on Zenodo (https://zenodo.org/records/20204118).

## Acknowledgements

We thank Adam Dowle and Chloë Baldreki from the York Bioscience Technology Facility Metabolomics and Proteomics group for expert advice, TimsTOF runs and protein searches. We thank Karen Hogg, Graeme Park and Sukhveer Mann from the York Bioscience Technology Facility Flow Cytometry Core for expert advice, training, flow cytometer maintenance and QC. We thank Grant Calder from the York Bioscience Technology Facility Imaging Core for expert advice, training and microscope maintenance and QC. We thank Alastair Droop and Matthew Care from the York Bioscience Technology Facility Data Science Hub for code review and expert advice. We thank Mark Bentley in the Department of Biology Mechanical Workshop for custom manufacturing of tube holders for proteomic sorts and processing. We also thank all umbilical cord blood donors and their families.

WG acknowledges funding from: Leukaemia U.K. (2021/JGF/002), The Children’s Cancer and Leukaemia Group (CCLG 2022 13 Grey), The Medical Research Council (MR/X007146/1, National Mouse Genetics Network Director’s Fund MC_PC_21048 & Discovery Medicine North PhD studentship), The National Centre for the Replacement, Refinement and Reduction of Animals in Research (UKRI630), The European Haematology Association (RG112), The British Society for Haematology.

DH acknowledges funding from: Anthony Nolan and The David and Ruth Lewis Foundation. Special thanks to all the cord blood donors who consented for research use and the Anthony Nolan Cord Blood Programme.

The York Centre of Excellence in Mass Spectrometry was created thanks to a major capital investment through Science City York, supported by Yorkshire Forward with funds from the Northern Way Initiative, and subsequent support from EPSRC (EP/K039660/1; EP/M028127/1).

## Author contributions

Conceptualization: ZM, WG. Performed Experimentation: ZM, LW, MG, WG. Data Acquisition: ZM, NS, LW, MG, WG. Analyzed Data: ZM, NS, MG, DMD, WG. Writing (original draft): ZM. Writing (reviewing, additions and editing): ZM, NS, LW, MG, DMD, DH, WG. Funding Acquisition: DH, WG. Project Administration: WG. Supervision: DH, WG.

## Competing interest declaration

The authors have no conflict of interest.

## Additional information

Supplementary information is available for this paper.

Correspondence and requests for materials should be addressed to William Grey.

## Notes

https://zenodo.org/records/20204118

### Competing Interest Statement

The authors have declared no competing interest.

