## Supplementary Information for "Spatiotemporal Proteome Remodeling Directs Human Hematopoietic Stem Cells via a Metabolic-Cytoskeletal Axis"

### Supplementary Figures

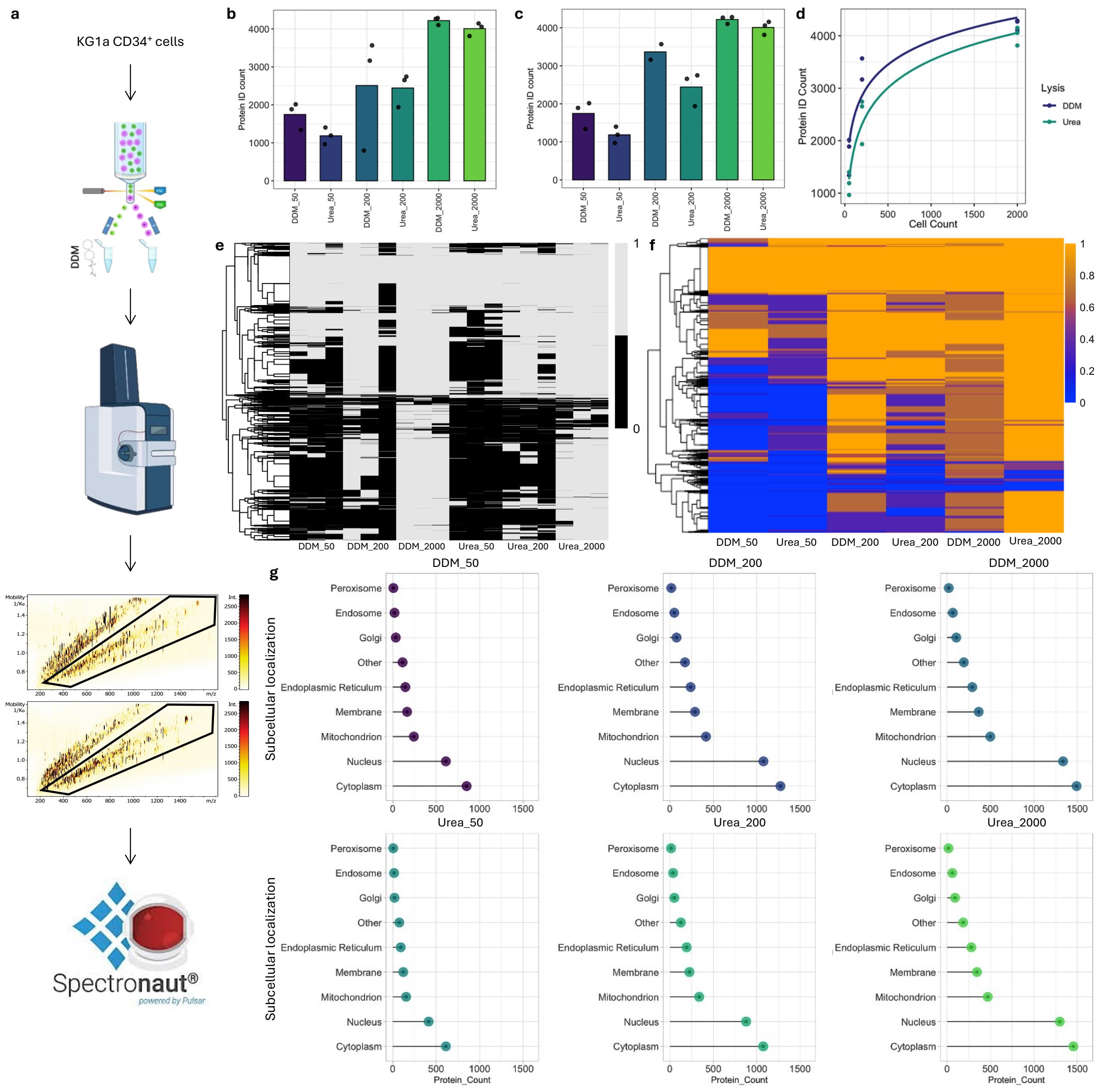


**Supplementary Figure 1. Benchmarking a robust ultra-low input proteomics pipeline using TimsTOF technology. a**, Schematic representation of the development of a low-input proteomics pipeline using the KG1a cell line and TimsTOF technology, **b,** Total number of all proteins identified from 50, 200 and 2000 cells using DDM or Urea lysis protocols, **c,** Total number of proteins identified from minimum 2 peptides and FDR<0.01, excluding one misaligned DDM-sorted sample, from 50, 200 and 2000 cells using DDM or Urea lysis protocols, **d,** Overlay of the relationship between number of input cells and count of identified proteins from DDM vs urea lysis, **e**, Heatmap of missing values from every sample for 50, 200 and 2000 cells using DDM or urea lysis protocols, **f**, Heatmap of data completion per cell count input for each lysis buffer to indicate overall coverage (1 equal complete detection, 0 equals not detected), **g**, Count of proteins across subcellular localizations for each cell input and analysis condition to show subcellular depth of detection.

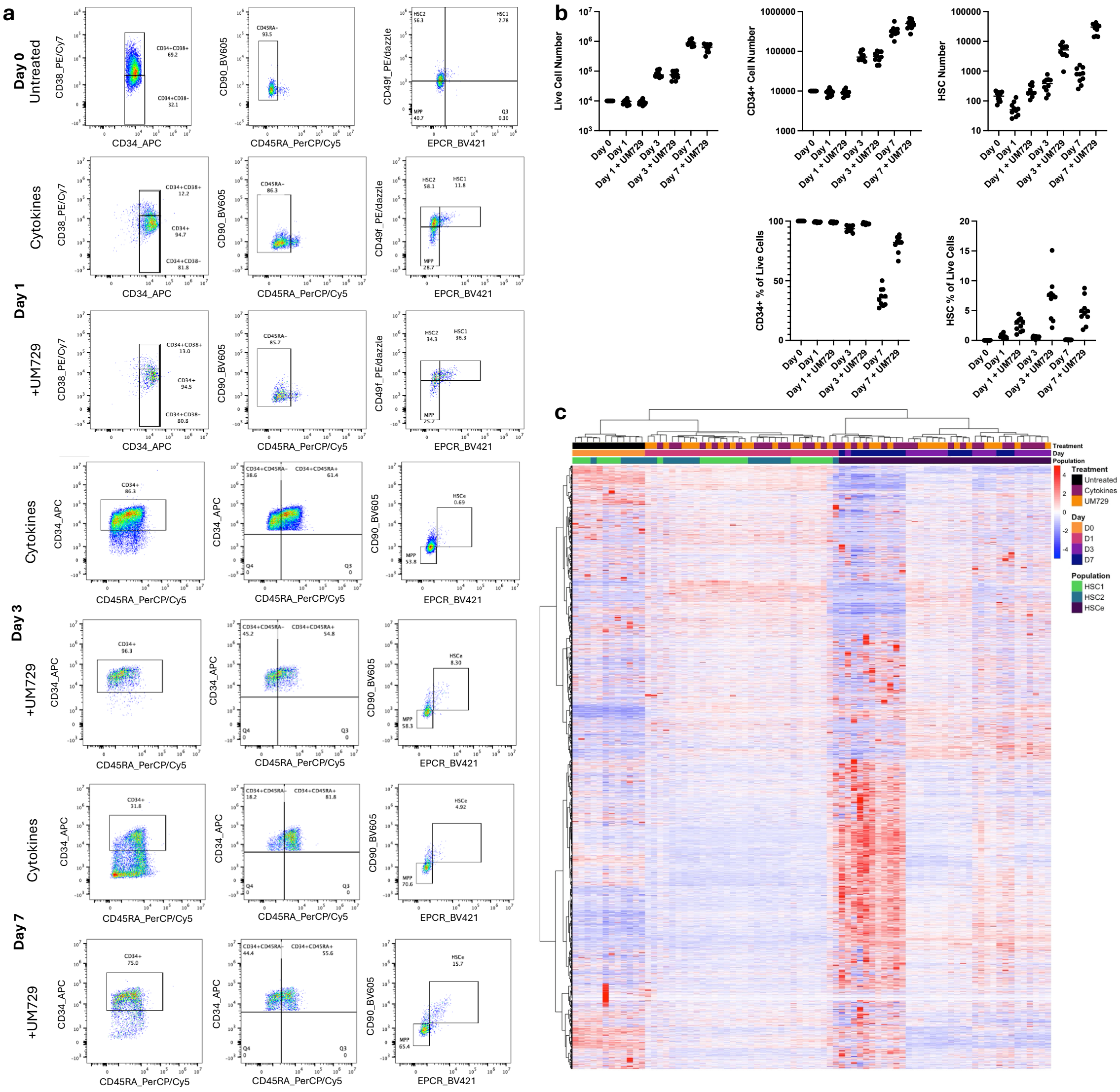

**Supplementary Figure 2. CB HSPC flow cytometry sorting, analysis strategy, and culture outcomes. a**, Flow cytometry gating and sorting strategy for HSC$^1$, HSC$^2$ and HSC$^e$ from *de novo* day 0 (untreated) cells or cells cultured in basal media (cytokines) or with UM729 for 1, 3 or 7 days, **b,** Absolute count of total, CD34$^+$, and phenotypic HSCs (HSC$^1$, HSC$^2$ or HSC$^e$) as well as % in live cells for 0, 1, 3 or 7 days in culture, **c**, Heatmap of all proteins detected showing clustering across all untreated or cultured HSCs.

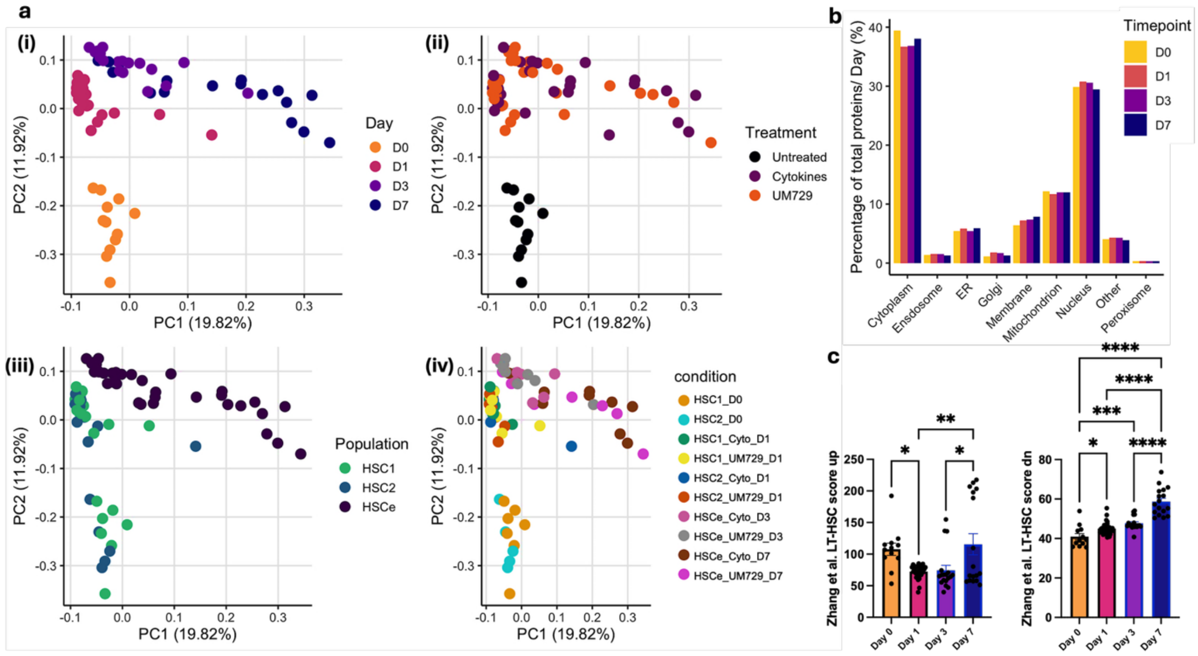


**Supplementary Figure 3. a,** PCA plots based on differentially abundant proteins in total HSC proteomics across days (i), treatment (ii), HSC type (iii) or combination of all conditions (iv). **b,** Percentage of proteins identified in each subcellular compartments within the detected proteome on each day, **c,** Bar chart of long-term HSC up- or down-regulated scores based on Zhang et al. (2022).

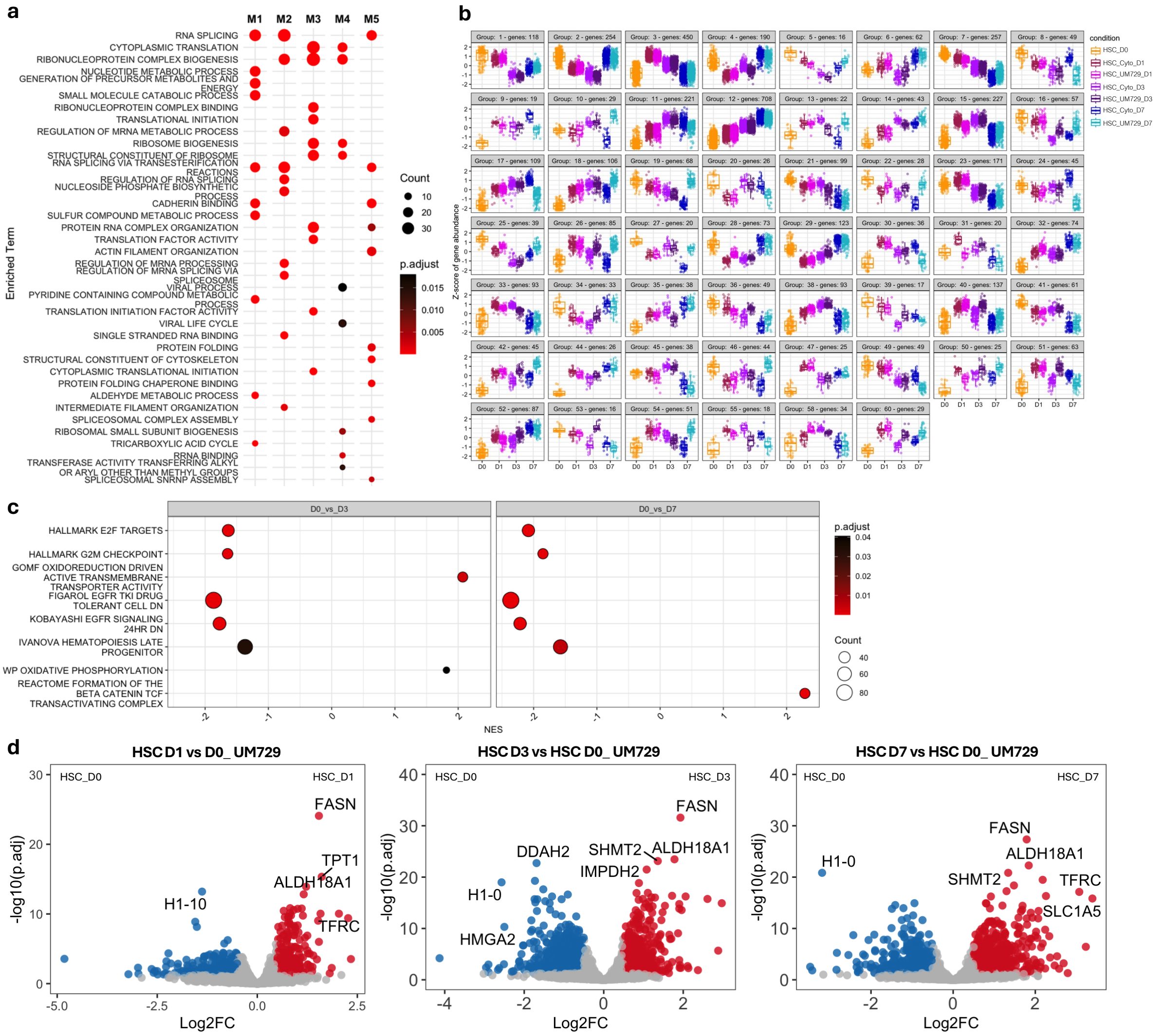

**Supplementary Figure 4. Overview of gene ontology and temporal changes in protein abundance patterns in HSCs. a,** Gene ontology enrichment analysis of the 5 Tradeseq modules generated by pseudotime comparison in HSCs, **b**, All groups of protein abundance changes from DEGpatterns across untreated and cultured HSCs ranging from 16 to 450 proteins/group, **c**, Pathways from GSEA analysis of overall DE proteins in D0 vs D3 or D0 vs D7 HSCs, **d,** Volcano plot showing differences in protein abundance in HSCs across all days of culture supplemented with UM729.

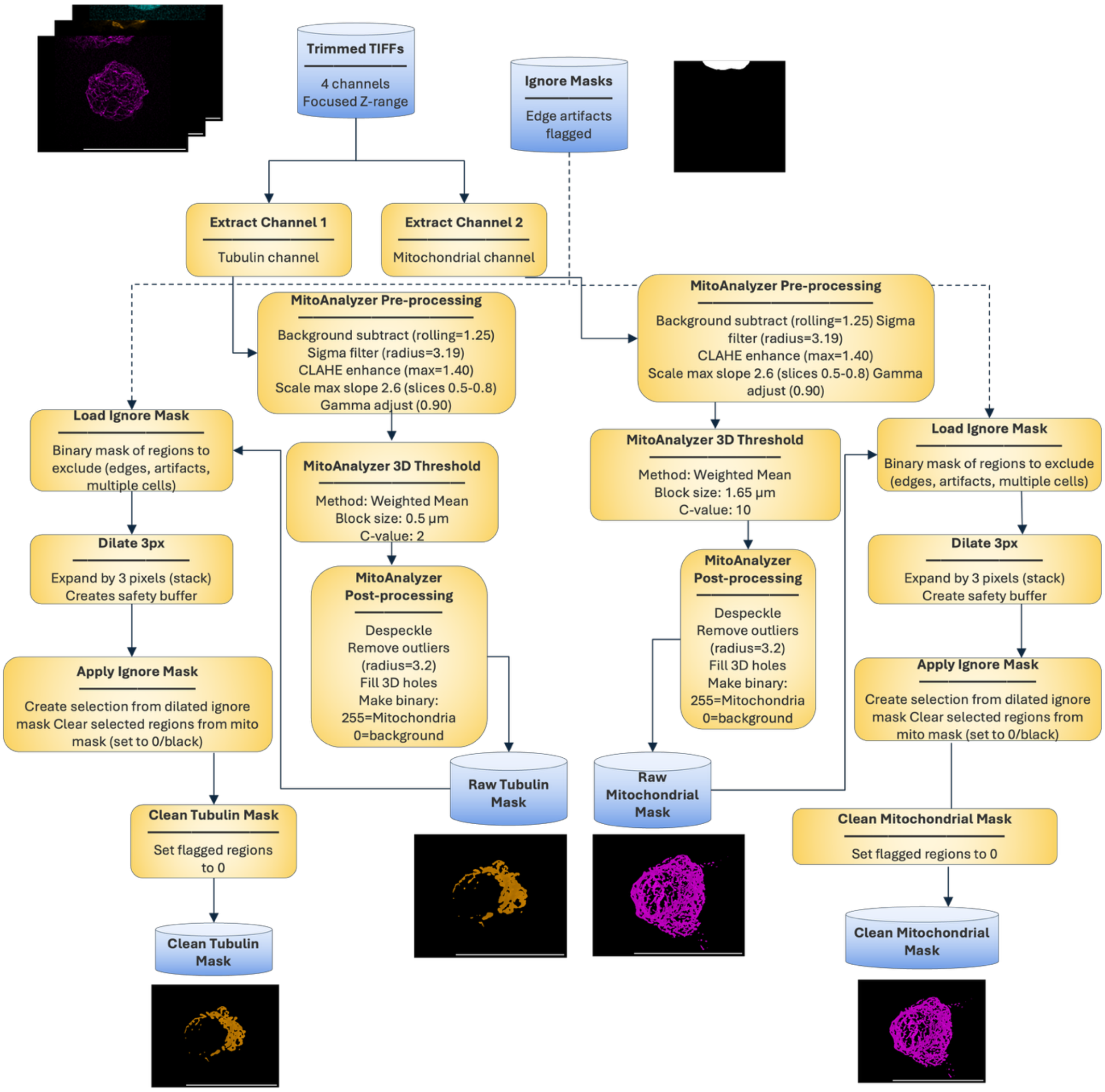


**Supplementary Figure 5. Representation of microscopy analysis workflow.** Overall analysis pipeline and representative masks generated for quantitative analysis of the mitochondrial and cytoskeletal structures and networks.

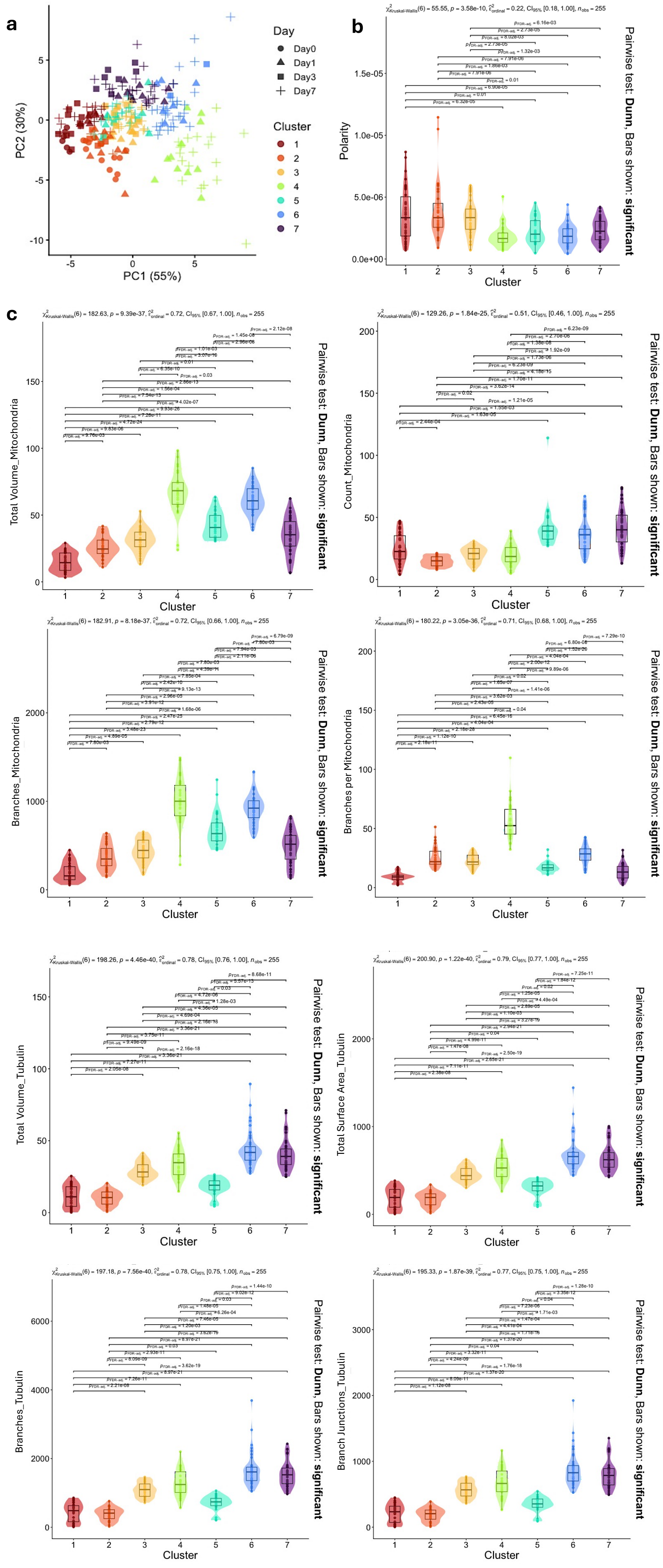


**Supplementary Figure 6. Analysis of cytoskeletal and mitochondrial features using unsupervised clustering.**

**a**, Unsupervised clusters based on morphological features presented on PCA plot, **b,** Violin plots showing values and statistical comparison of polarity across clusters (significant differences and p values shown in graph), **c,** Violin plots showing values and statistical comparison of tubulin and mitochondrial features including volume (μm$^3$), surface area (μm$^2$), etc. across clusters (significant differences and p values shown in graph).

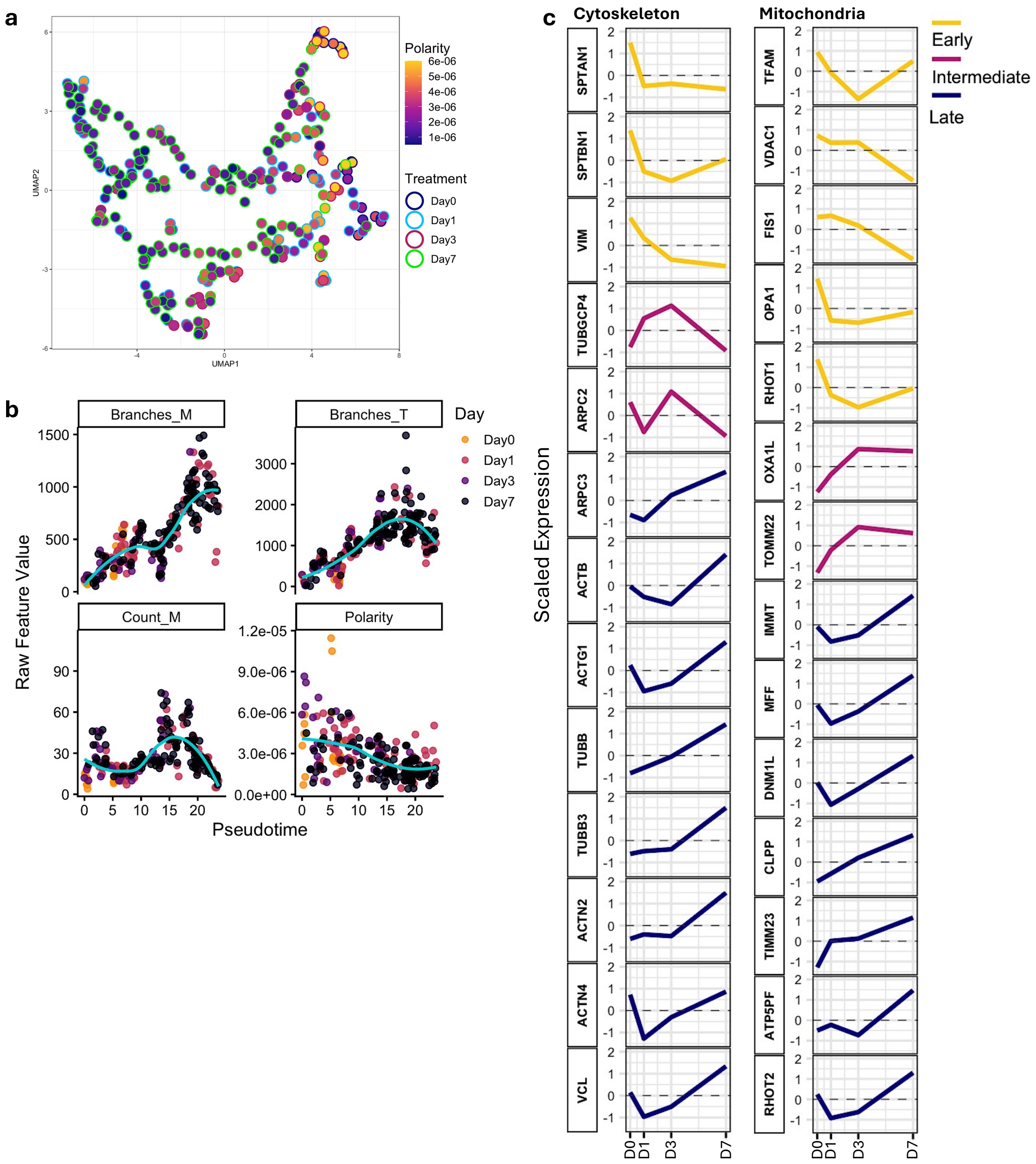


**Supplementary Figure 7. Trajectories of morphological changes across pseudotime and proteomic dynamics across days in culture. a**, Overlay of polarity values on UMAP with data point edges are colored based on days in culture, **b,** Tubulin branches, mitochondrial branches and counts and polarity trajectories across pseudotime in untreated samples colored by day in culture, **c,** Line trajectory of scaled expression of proteins associated with cytoskeletal structure as well as mitochondrial biogenesis, fission and fusion, and ATP production/metabolism from total proteomics of HSCs across days in culture.

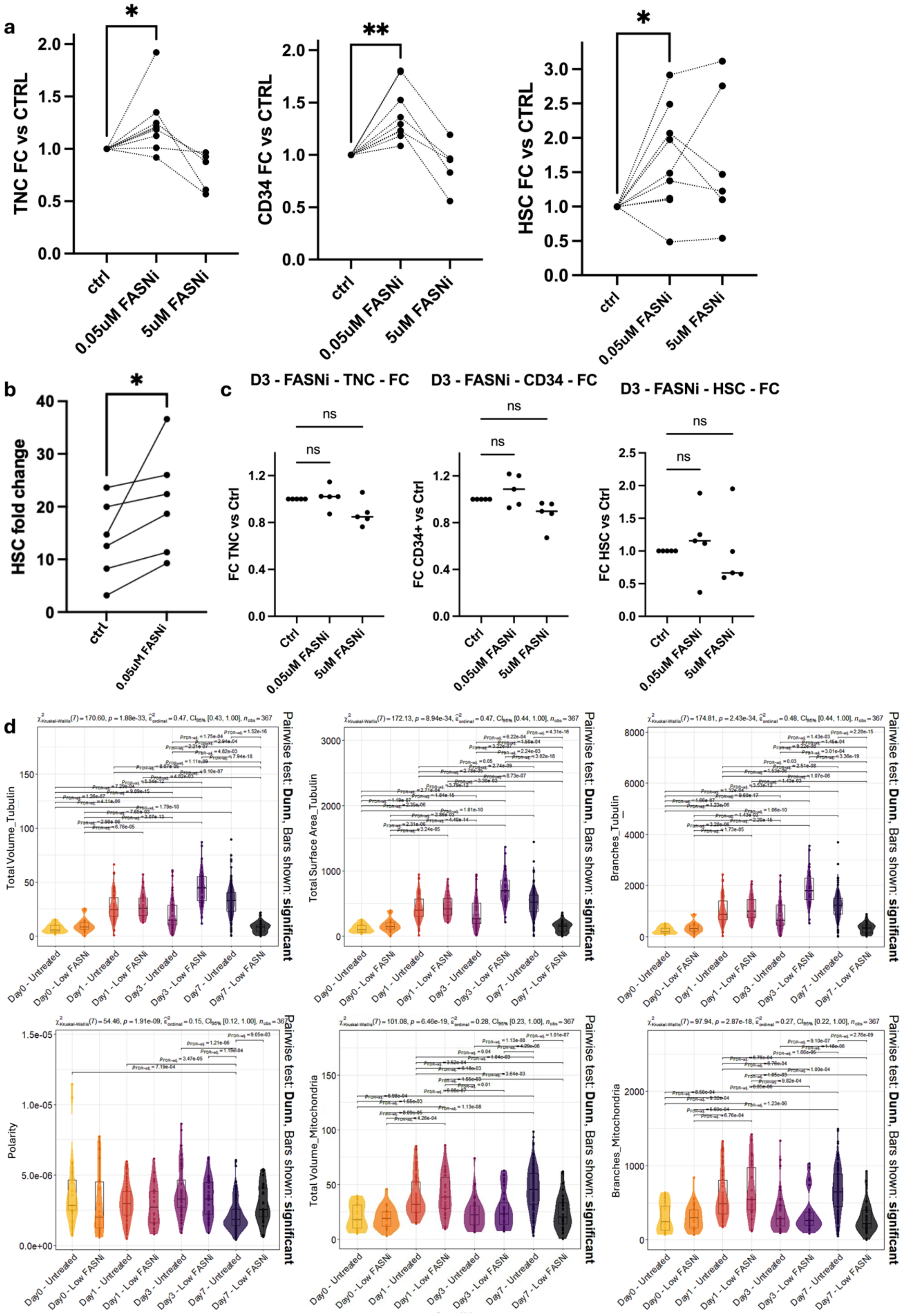


**Supplementary Figure 8. a,** Fold change in the total nucleated cells, CD34$^+$ cells and phenotypic HSCs in FASN-inhibited vs untreated cells from treatment with TVB-2640 initiated on day 0 (one-way ANOVA was used to calculate significance, * = p<0.05, **

= p<0.005), **b**, Fold change vs input number of phenotypic HSCs (available from 6 of 9 samples) following FASN inhibition initiated on day 0, **c,** Fold change vs control in the number of total nucleated, $CD34^+$ cells, and phenotypic HSCs following FASN inhibition initiated on day 3 of culture, **d,** Violin plots of values and comparative statistical analysis of temporal changes in morphological features including polarity in untreated and FASNi-treated cells (significant differences and p values shown in graph).

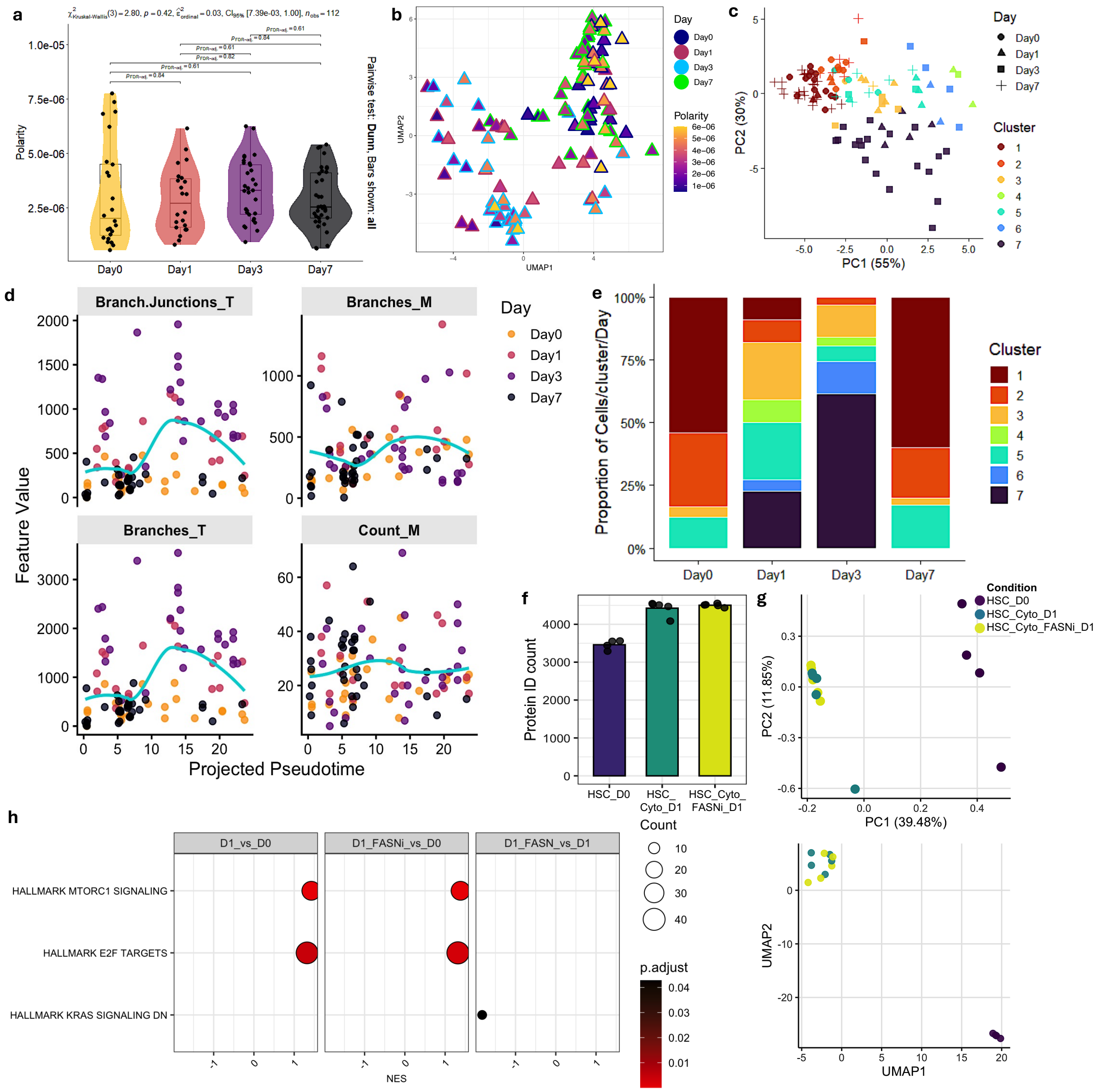

**Supplementary Figure 9. FASN activity is central to HSC adaptation to culture and its inhibition changes HSC expansion trajectory and adaptation. a,** Violin plots of polarity value and statistical comparison across FASNi-treated cells for each day (all p.adj values shown in graph), **b,** Polarity overlay of FASNi cells on the pseudotime trajectory, showing higher polarity in cells in later days in culture as distinguished by the edge color, **c,** Overlay of mcluster-based clustering of FASNi cells on PCA plots, **d,** Trajectories of morphological changes in FASNi-treated cells over pseudotime indicating a distinct drop in mitochondria branching in later pseudotime in FASNi cells in contrast to the rise observed in untreated cells, **e**, Proportion of cells from each morphological cluster comprising total cell population on each day following FASNi treatment, **f**, Bar chart of number of identified proteins in FASN inhibition experiment, **g**, PCA plot and UMAPs from proteomics outcome, **h**, GSEA analysis indicating changes in mTORC2 and E2F and RAS signatures in day 1 FASNi cells compared to D0 or D1 untreated cells.

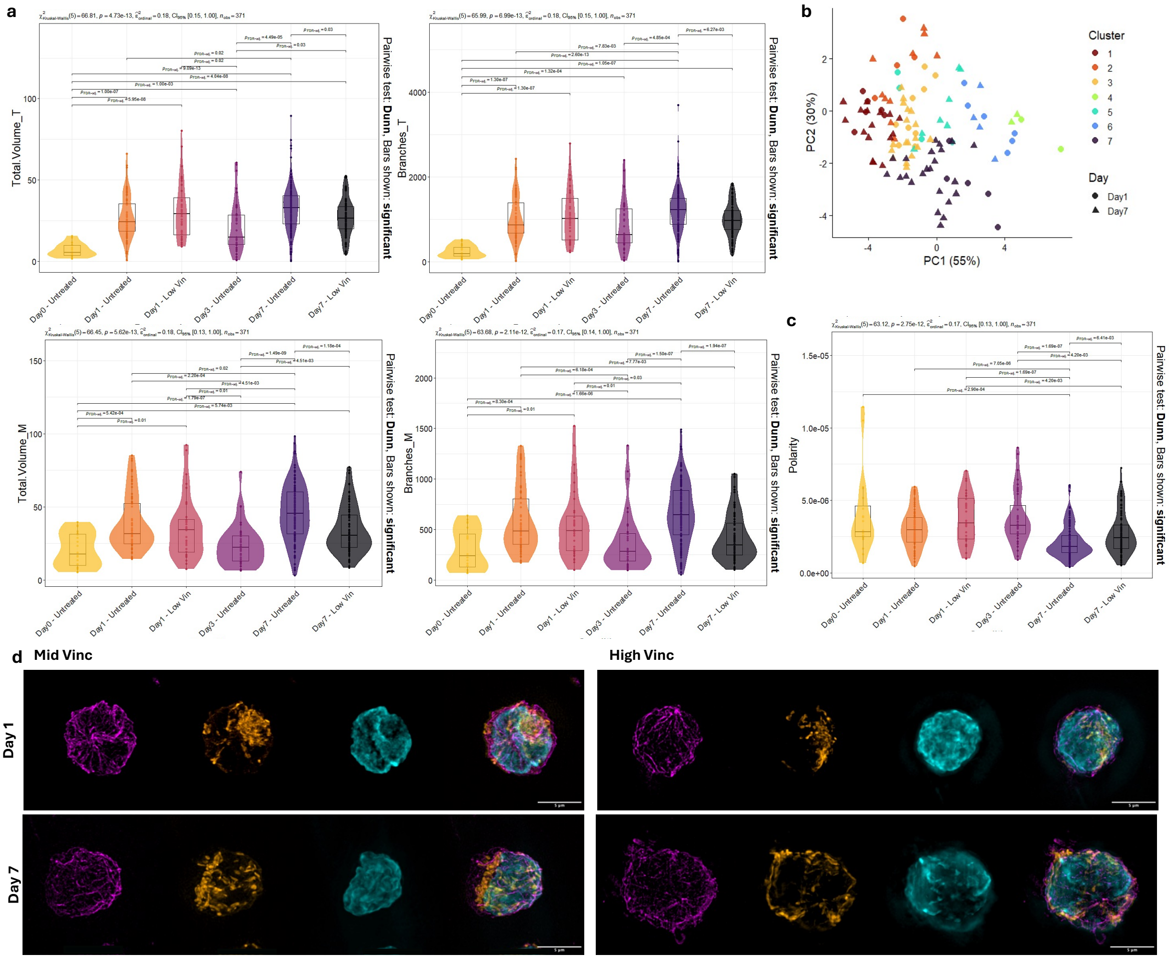


**Supplementary Figure 10. Vincristine-based disruption of cytoskeletal scaffolding derails structural and metabolic organization of HSC adaptation. a,** Violin plots of values and comparative statistical analysis of temporal changes in cytoskeletal and mitochondrial features in low dose vincristine-treated vs untreated cells (significant differences and p values shown in graph) (T=tubulin, M=mitochondria), **b**, Overlay of mcluster-based clustering of low dose vincristine-treated cells on PCA plots, **c,** Violin plots of values and comparative statistical analysis of temporal changes in polarity in vincristine-treated vs untreated cells (significant differences and p values shown in graph), **d**, Representative microscopy images of cells treated with medium or high dose of vincristine indicating morphological changes in tubulin (magenta), mitochondrial features and polarity (MitoTracker Red, orange), nucleus (DAPI, cyan) and overlay of all channels across days in culture (scale bar is 5μm).
